# *H19* isoform switching during neuronal differentiation reveals a distinct pluripotency-associated non-canonical transcript

**DOI:** 10.64898/2026.11.25.769322

**Authors:** Nicholas J. Guzman, Luisa M. Abadia, Yanelee L. Felicien, Michael J. Woodburn, Srinivas Somarowthu, Ridhdhi Desai, Christian Sell

## Abstract

*H19* is one of the earliest discovered and most extensively studied lncRNAs, yet most studies have focused on the canonical 2,315-nt transcript arising from the *IGF2/H19* locus on chromosome 11. We examined *H19* isoform usage during the pluripotency-to-neuronal transition using NGN2/ASCL1-induced neurogenesis in KOLF2.1J human induced pluripotent stem cells (hiPSCs). We show that a smaller, non-canonical isoform, variant 3, is the predominant *H19* isoform in the pluripotent cells examined. This isoform lacks the large canonical promoter-derived exon 1 and contains a short, unique first exon encoded within the imprinting control region upstream of *H19*. Upon neuronal induction, isoform-specific RT-qPCR showed that variant 3 declined below the lower limit of reliable quantification (LLRQ) as canonical *H19* and *IGF2* mRNA increased. In contrast, human bone marrow-derived mesenchymal stem cells expressed canonical *H19*, while variant 3 remained below the LLRQ during neurogenic, osteogenic, and adipogenic differentiation. Independent FANTOM5 CAGE datasets supported transcription initiation in the predicted variant 3 promoter region in hiPSCs and human embryonic stem cells. Long-read RNA-seq identified transcripts linking the variant 3 first exon to downstream *H19* exons. Public OCT4 and SOX2 ChIP-seq datasets showed recurrent peak overlap at the upstream OSM1 candidate region in pluripotent cells, with corresponding peaks absent in the differentiated datasets examined. Together, these findings support cell-state-associated regulation of the *IGF2/H19* locus and identify alternative *H19* isoform usage as a feature of human pluripotent cells.

## Introduction

Long non-coding RNA (lncRNA) loci frequently produce multiple transcript isoforms through both alternative transcription start site (TSS) usage and alternative splicing ^1–3^. These distinct isoforms can differ substantially in sequence leading to differences in structure, localization, and molecular interactions resulting in distinct regulatory and biological functions. These processes are often cell-state dependent; for example, a differentiation-driven shift in TSS usage was reported for *HOTAIR* during adipogenic differentiation^4^, and changes in transcript initiation, exon usage, and polyadenylation site selection have also been identified during neuronal differentiation^5^. However, lncRNAs have historically been studied primarily at the gene level, and many commonly used assays target regions shared among multiple transcripts. Therefore, cell-state-dependent changes in isoform usage may be interpreted simply as changes in total gene expression, while the expression dynamics and functional differences of individual transcripts remain unresolved.

*H19* was one of the earliest lncRNAs identified and remains one of the most extensively studied^6–8^. The *H19/IGF2* locus was also among the first discovered imprinted gene loci and is highly conserved in therian mammals^9–11^. Parent-of-origin-specific methylation results in a hypomethylated imprinting control region (ICR1) on the maternal allele and a hypermethylated ICR1 on the paternal allele^12^. This allows selective binding of the insulator protein CTCF to maternal ICR1 and establishes a higher-order chromatin conformation that restricts downstream enhancers from activating *IGF2* and instead favors *H19* expression^13–16^. On the hypermethylated paternal allele, CTCF binding is blocked and those same enhancers can drive *IGF2* expression. Additionally, conserved Sox-Oct motifs are present within ICR1. In mouse embryonic carcinoma cells and developmental models, the pluripotency factors OCT4 and SOX2 bind these motifs and contribute to maintenance of ICR1 hypomethylation^17^. Disruption of ICR1 methylation and reciprocal *H19/IGF2* expression contribute to the opposing growth phenotypes observed in Beckwith-Wiedemann and Silver-Russell syndromes^18–20^. Most work involving *H19* has focused on the 2,315-nt canonical transcript, which contains miR-675 within its large first exon^21,22^. *H19* can also function through binding to RNA-binding proteins, including chromatin-modifying complexes, and through sequestration of miRNAs as a competing endogenous RNA, underscoring its context-specific roles in embryonic and adult tissues^8^.

*H19* is highly expressed during embryonic development and is generally downregulated after birth, although expression is retained or reactivated in specific adult, stem-cell, and disease contexts^8^. *H19*-derived miR-675 contributes to adult hematopoietic stem cell quiescence through restriction of IGF1R signaling^23^. During osteogenic differentiation in several stem-cell models, *H19* has been reported to be upregulated and to promote differentiation through mechanisms including miR-675/TGF-beta1/Smad3/HDAC and *H19*/FOXC2/Wnt/beta-catenin signaling^24–26^. *H19* is also re-expressed in numerous cancers and has been assigned both oncogenic and tumor-suppressive functions depending on the cellular context^27–30^.

*H19* and *IGF2* have also been implicated in the pluripotent state and neural differentiation. In pluripotent human embryonic carcinoma and embryonic stem cells (hESCs), knockdown of *H19* decreased expression of pluripotency factors and produced molecular changes consistent with early differentiation^31^. In the developing mouse, *H19* and *Igf2* are expressed in specific neuroectoderm-derived regions of the brain and spinal cord^32^, and activation of the chromosome 11p15.5 region containing *H19* and *IGF2* has been observed during the production of neuronal progenitor cells from hESCs^33^. IGF2 has additionally been shown to regulate adult neural stem-cell proliferation and terminal differentiation^34,35^. In one induced neural stem-cell model, *H19* expression decreased during the transition from induced pluripotent stem cells (iPSCs) to induced neural stem cells and neurons, while *H19* knockdown promoted neural differentiation^36^. Together, these studies establish that *H19*/*IGF2* expression changes during pluripotency and neural differentiation, but they do not resolve which *H19* transcript isoforms are expressed in these different cellular states. The apparently divergent functions assigned to *H19* may reflect genuine cell-type-and stage-dependent functions, but they may also partly result from assays that do not distinguish transcripts with different sequences and potentially different functions.

Although *H19* has generally been studied at the gene or canonical-transcript level, seven *H19* transcript variants are currently annotated by NCBI RefSeq^8,21^. Six variants share the large 1,325-nt canonical exon 1 and exhibit comparatively minor differences among their downstream exons. In contrast, the 1,039-nt non-canonical transcript annotated as variant 3 entirely lacks the canonical exon 1 and instead contains a 49-nt alternative exon 1 followed by the same four downstream exons present in canonical variant 1. This unique exon is encoded within the upstream ICR1 regulatory region that also contains conserved Sox-Oct motifs^17,21^. Variant 3 is therefore less than half the length of canonical *H19* and cannot serve as a precursor for miR-675, which is encoded within the canonical exon 1^21,22^. A recent study identified a 131-amino-acid open reading frame beginning within the variant 3-specific first exon that encodes the microprotein GMRSP in human vascular smooth muscle cells^37^. Thus, although *H19* is conventionally classified as a lncRNA, variant 3 also has demonstrated coding potential in that cellular context. Its distinct first-exon architecture is consistent with generation through alternative promoter or TSS usage rather than through alternative splicing of the canonical *H19* precursor.

Here, we investigated *H19* isoform usage during the pluripotency-to-neuronal transition using a rapid and tractable, doxycycline-inducible NGN2/ASCL1 neurogenesis system in KOLF2.1J human iPSCs^38–40^. Using calibrated isoform-junction reverse transcription-quantitative polymerase chain reaction (RT-qPCR), we identified variant 3 as the predominant detected *H19* isoform in pluripotent iPSCs and hESCs. Variant 3 rapidly declined below lower limit of reliable quantification (LLRQ) during exit from pluripotency, coinciding with transient induction of canonical *H19* and *IGF2* during early loss of pluripotency/neuronal differentiation. In contrast, adult bone marrow-derived mesenchymal stem cells (BMSCs) expressed canonical *H19* while variant 3 remained below LLRQ before and during differentiation into multiple lineages. Additionally, variant 3 was not reliably quantifiable in the adult or cancer-derived cell types examined.

FANTOM5 cap analysis of gene expression (CAGE) data support this pluripotency-associated alternative initiation region and long-read RNA-seq confirms expected splicing to downstream exons, while independent OCT4 and SOX2 ChIP-seq datasets demonstrate recurrent peak overlap at nearby candidate regions in pluripotent cells and absence of corresponding formal peak calls in the differentiated comparator tracks examined. Together, these findings identify a cell-state-dependent transition in *H19* isoform usage and support variant 3 as a pluripotency-associated non-canonical *H19* transcript. To our knowledge, this is the first calibrated, isoform-junction-resolved demonstration that variant 3 is the predominant detected *H19* isoform in human pluripotent stem cells and that dominant *H19* isoform usage changes during lineage specific differentiation.

## Results

### NGN2/ASCL1 induction rapidly reduces pluripotency marker expression and initiates neuronal differentiation in KOLF2.1J hiPSCs

To define the expression dynamics of transcript isoforms within the *H19/IGF2* locus during early neurogenesis, we transduced a doxycycline-inducible NGN2/ASCL1 plasmid^40^ into KOLF2.1J iPSC cells by lentivirus. This rapid and tractable system produces induced neuron-like cells (iNeurons). We first wanted to validate the rapid loss of pluripotency and induction of neuronal programs upon induction with doxycycline (Fig. 1A). The cells rapidly adopt a neuron-like morphology following induction, with increased neurite extension and branching throughout 14 days of maturation (Fig. 1B). RT-qPCR showed that *POU5F1* (encoding OCT4) and *NANOG* are rapidly decreased (Fig. 1C). *SOX2* initially decreased at day 1, followed by an increase compared to day 1 at days 2 and 3 (still significantly decreased from baseline), and decreased again at day 7; this transient *SOX2* expression is consistent with its established role in maintaining neural progenitor identity^41^ (Fig. 1C). As well, *MAP2* is strongly increased by day 7 (Fig. 1C). Consistent with the transcriptional profile, immunofluorescence showed robust expression of βIII-tubulin and MAP2 in day 10 iNeurons (Fig. 1D). These data confirm that our KOLF2.1J iNeuron model, derived from the 8^th^ clone tested (and subsequently named KOLF2.1J-C8), produces a rapid and tractable pluripotency-to-neuronal transition that can be used to explore isoform dynamics and regulation of the *H19/IGF2* locus.

**Figure 1.**
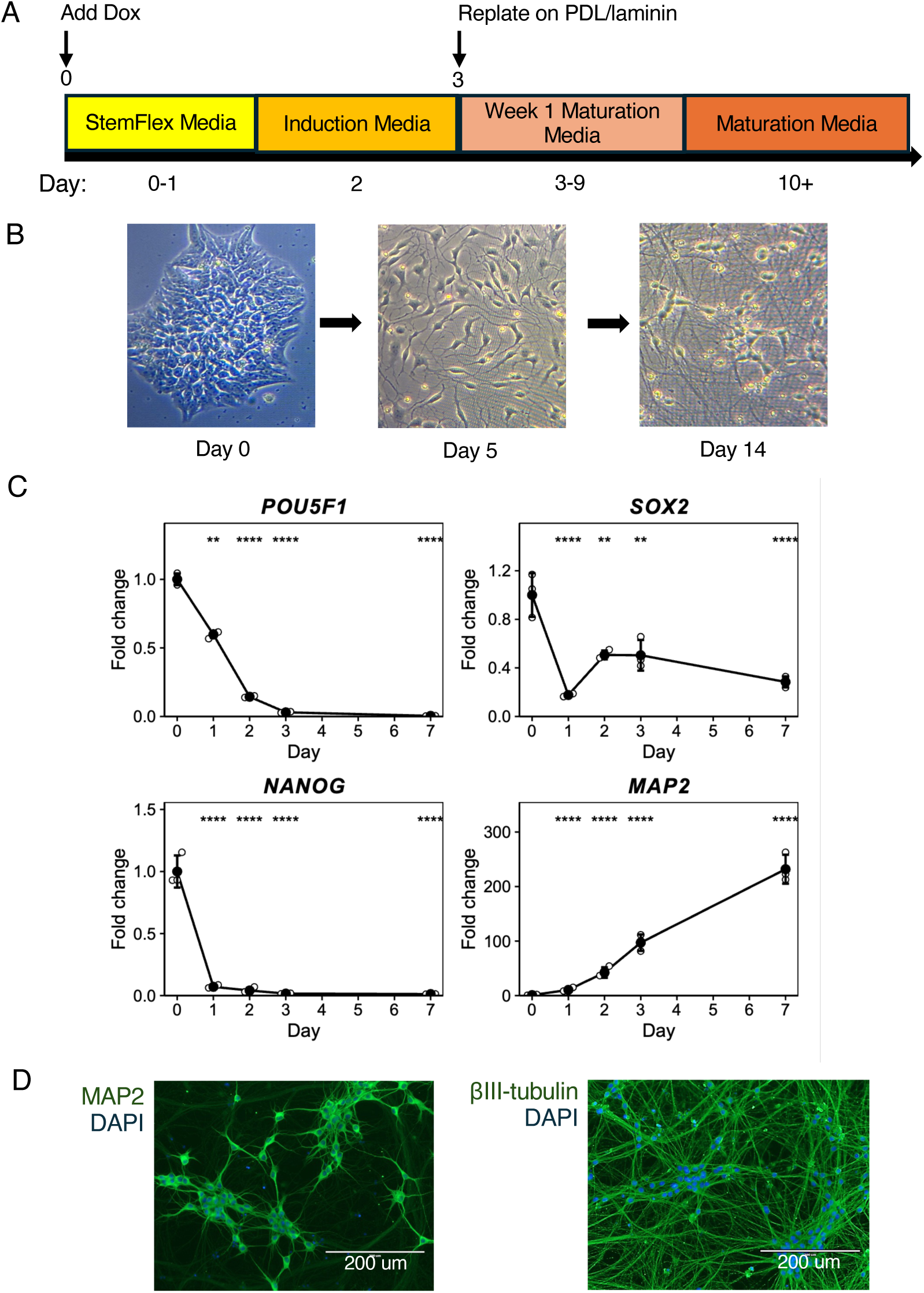
NGN2/ASCL1-induced neurogenesis in KOLF2.1J-C8 hiPSCs. **(A)** Schematic of doxycycline induction and iNeuron maturation. **(B)** Representative phase-contrast images at days 0, 5, and 14. **(C)** 18S-normalized RT-qPCR of *POU5F1* (encoding OCT4), *SOX2*, *NANOG*, and *MAP2* relative to day 0. Filled points indicate geometric means, open points individual biological samples, and error bars SD of biological-sample fold changes (n=3 per time point). Comparisons versus day 0 used one-way ANOVA on ΔC_T_ values with Dunnett-adjusted contrasts (**P<0.01, ***P<0.001, ****P<0.0001). **(D)** Representative day-10 iNeuron immunofluorescence for MAP2 or βIII-tubulin (green) and DAPI (blue); scale bars as indicated.

### *H19* variant 3 predominates during pluripotency and canonical *H19* predominates during differentiation

As *H19* has seven annotated transcript isoforms in NCBI RefSeq^21^, we wanted to determine *H19* isoform levels during the pluripotent-to-neuronal transition. Of the seven annotated transcripts (Fig. 2A), six share the large canonical exon 1 (further referred to as ‘canonical isoforms’) while variant 3 contains a short alternative first exon contained within the ICR1^17,21,42,43^ (Fig. 2A and Fig S1A). Primer sets were designed to amplify unique exon-exon junctions of the seven isoforms, with some isoforms sharing splice sites (Fig. 2A and Fig. S1B). Preliminary SYBR Green RT-qPCR showed increased signal for the canonical-isoform primer sets but decreased signal for the variant 3 primer set from undifferentiated KOLF2.1J-C8 cells to day 3 iNeurons (Fig. 2B). Sequencing of the gel-purified and reamplified RT-qPCR product from the undifferentiated KOLF2.1J-C8 cells confirmed a match to *H19* variant 3 (Fig. S1D). We therefore optimized assays targeting the distinct exon 1-2 junctions of canonical *H19* and variant 3 (Fig. S1B). *H19* variant 1 (canonical) and variant 3 were cloned into in-vitro transcription (IVT) vectors, and the resulting RNAs were serially diluted to generate standard curves spanning 1,000-10,000,000 copies/reaction. This range approximates 0.5-5,000 copies per cell equivalent (assuming ∼0.01 ng RNA per cell) and was used to validate the SYBR Green assays. Optimized primers showed linear amplification across the RNA input dilutions (Fig. 2C). *H19* assay signals were classified as below the lower limit of reliable quantification when the expected melt peak was absent or did not dominate the nonspecific peaks in any technical replicate (Fig. S3A). Each assay produced a defined specific melt peak (88°C for canonical *H19* and 86°C for variant 3), and neither assay produced the expected specific melt peak with the noncognate template (Fig. S1C). These data demonstrate that this assay reliably distinguishes canonical and non-canonical *H19* isoforms.

**Figure 2.**
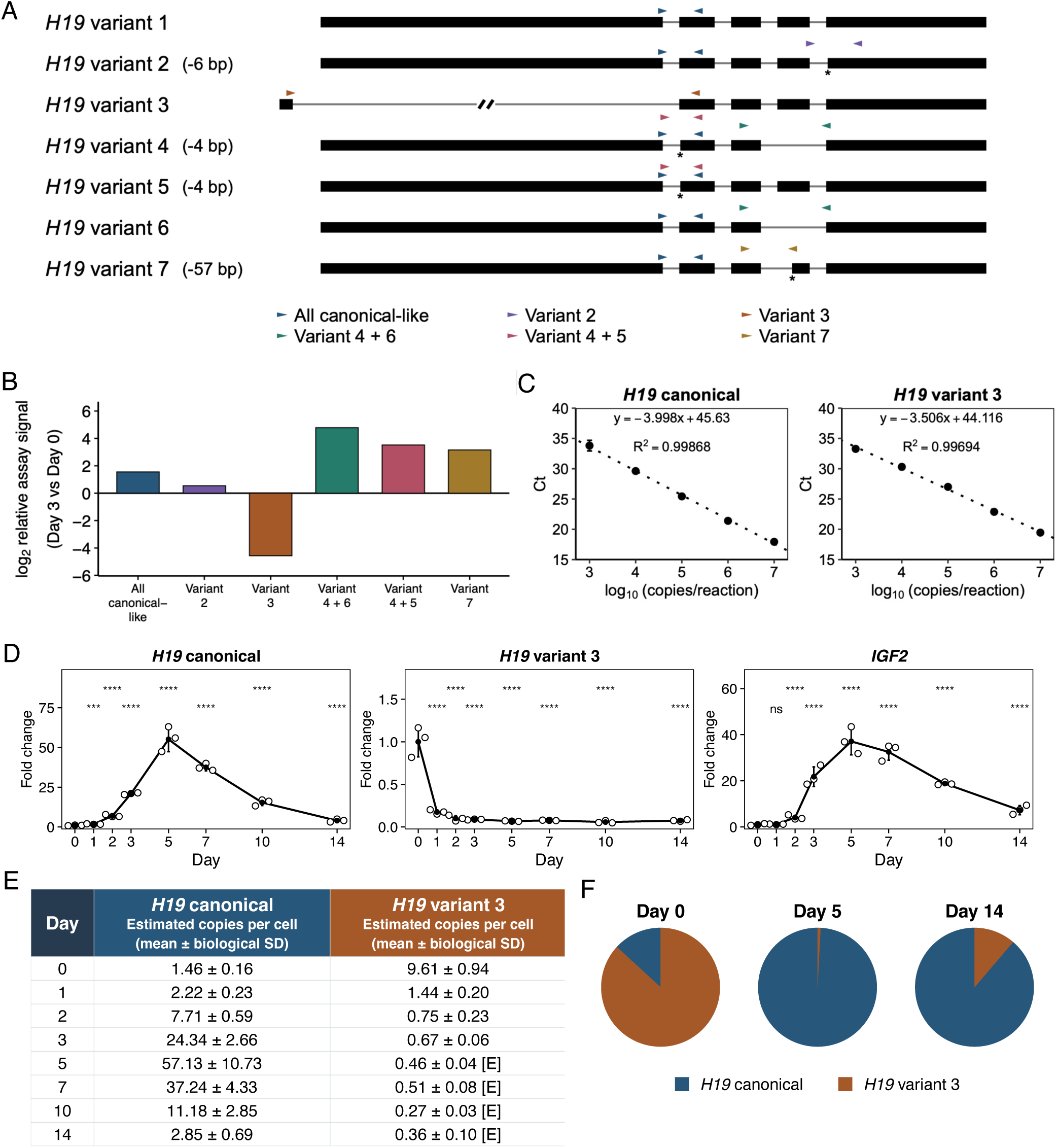
***H19* isoform switching during NGN2/ASCL1-induced neurogenesis. (A)** Seven NCBI RefSeq-annotated *H19* transcript variants and RT-qPCR assay positions; parenthetical values indicate exon shortening in base pairs, and asterisks mark affected exon boundaries. **(B)** Preliminary SYBR Green RT-qPCR screen of *H19* transcript assays, expressed as log₂ relative assay signal at day 3 versus day 0 (C_T_ [day 0] – C_T_[day 3]), with one biological sample per condition. (C) Standard curves generated from in vitro-transcribed (IVT) canonical *H19* and variant 3 RNA (10³–10⁷ copies/reaction; technical duplicates). Points indicate means and error bars technical SD; regression equations and R² are shown. **(D)** Efficiency-adjusted, 18S-normalized RT-qPCR of canonical *H19*, *H19* variant 3, and *IGF2* during iNeuron differentiation, relative to day 0. Filled points indicate geometric means, open points individual biological samples, and error bars SD of biological-sample fold changes (n=3 per time point). Comparisons versus day 0 used one-way ANOVA on log₂-transformed normalized quantities with Dunnett-adjusted contrasts (ns, not significant; ***P<0.001; ****P<0.0001). **(E)** IVT-standard-curve-derived estimated copies per cell equivalent (mean ± biological SD, n=3 per time point). Cell equivalents assume 20 ng RNA per reaction and 10 pg total RNA per cell. [E] marks time points with at least one biological-sample estimate outside the measured standard-curve range. **(F)** Estimated percentages of combined *H19* copies attributable to canonical *H19* and variant 3 at days 0, 5, and 14. Fractions were calculated within each biological sample and averaged across three biological samples. For **(D-F)**, values for samples classified below the lower limit of reliable quantification are exploratory assay-derived estimates.

Having established assays that distinguish canonical from non-canonical *H19*, we next profiled transcript dynamics during NGN2/ASCL1-induced neurogenesis. KOLF2.1J-C8 cells were differentiated for 14 days, with total cellular RNA collected across multiple time points to examine *H19* isoform dynamics during loss of pluripotency, early neurogenesis, and neuronal maturation. *IGF2* was additionally examined at each time point to assess its temporal relationship to *H19* isoform dynamics within the shared *H19/IGF2* locus. Canonical *H19* levels were below LLRQ at day 0 but increased sharply during neurogenesis, peaking at day 5 before declining toward baseline by day 14 (Fig. 2D and Fig. S2A). In sharp contrast, non-canonical variant 3 was expressed at a low but reliably quantifiable level at day 0 and rapidly fell below LLRQ as the cells exited pluripotency and entered neurogenesis (Fig. 2D and Fig. S2A). *IGF2* was detectable at day 0 and increased and subsequently decreased with a temporal profile similar to canonical *H19* (Fig. 2D). Copy estimates derived from RNA standard curves indicated that variant 3 predominated at day 0 whereas canonical *H19* predominated at days 5 and 14 (Fig. 2E-F). To determine whether variant 3 expression was unique to our reprogrammed iPSC model, we also assayed HuES8 hESCs and detected variant 3 while canonical *H19* remained below LLRQ (Figs. 3B and S2B). These data reveal that non-canonical *H19* variant 3 is the predominant detected *H19* isoform in the pluripotent states tested and that it declines below LLRQ while broader *H19/IGF2* locus activation occurs during loss of pluripotency and early neurogenesis.

**Figure 3.**
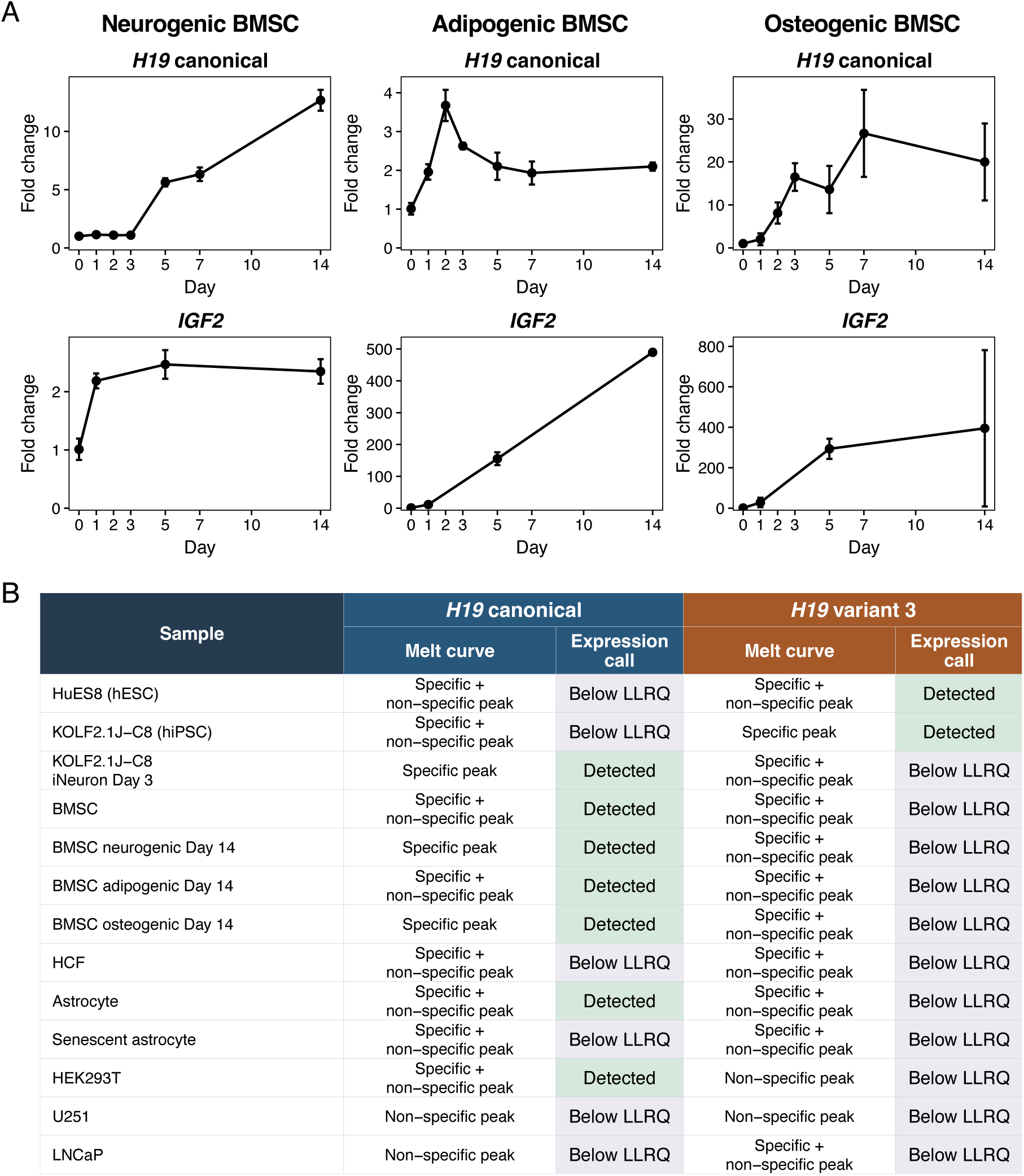
Canonical *H19* and *IGF2* expression during BMSC differentiation and *H19* isoform detection across additional cell types. **(A)** Efficiency-adjusted, 18S-normalized SYBR Green RT-qPCR of canonical *H19* and *IGF2* during neurogenic, adipogenic, and osteogenic differentiation of human bone marrow-derived mesenchymal stem cells (BMSCs), relative to the corresponding day-0 baseline. Each lineage was represented by one biological differentiation series. Points show means and error bars show SD across two to three quantifiable technical wells per target and time point. **(B)** Canonical *H19* and *H19* variant 3 melt-curve profiles and operational expression calls across the indicated cell types. LLRQ, lower limit of reliable quantification; classification criteria are illustrated in Fig. S3A.

### Canonical *H19* predominates in undifferentiated and differentiating BMSCs

We next asked whether adult multipotent stem cells also express *H19* variant 3. Unlike pluripotent cells, variant 3 was below LLRQ in undifferentiated adult bone marrow-derived mesenchymal stem cells (BMSCs), while canonical transcripts were detected at low abundance (Figs. 3B and S2C). BMSCs were then differentiated for 14 days toward neurogenic, osteogenic, and adipogenic lineages. Canonical *H19* increased during neurogenic and osteogenic differentiation and showed a smaller increase during adipogenic differentiation (Fig. 3A). *IGF2* was upregulated during osteogenic and adipogenic differentiation (Fig. 3A). Non-canonical variant 3 remained below LLRQ at baseline and during differentiation (Figs. 3B and S2C). We then examined additional cell types and found that, although canonical *H19* expression varied between cell types, variant 3 was not reliably quantifiable under the tested conditions (Figs. 3B and Fig. S3B-C). Thus, among the cell types and cells states assayed here, reliably quantifiable variant 3 signal was restricted to the pluripotent cell lines among the cells examined.

### RNAscope reveals relatively uniform *H19* signal in undifferentiated hiPSCs and heterogeneous signal in differentiating iNeurons and BMSCs

We next used RNAscope, an RNA-FISH-based method capable of visualizing individual RNA molecules, during differentiation of KOLF2.1J-C8 cells across the 14-day differentiation period (Fig. S4A, C). Because the tiled *H19* probe set targets sequence shared across *H19* transcripts, this assay cannot distinguish canonical and variant 3 isoforms and therefore provides qualitative context for the isoform-resolved RT-qPCR data. The probe revealed low but consistent signal across KOLF2.1J cells, which increased heterogeneously by day 5 and was lower by day 14, aligning with RT-qPCR results (Figs. S4A, C and 2D). In day 0 BMSC we observed very low signal for the majority of cells. Interestingly, the data revealed a small subpopulation of cells with much greater signal prior to in-vitro induced differentiation (Fig. S4B-C). At day 14 we observed a heterogeneous cell population demonstrating high and low probe signal, which is similar in heterogeneity to day 5 iNeurons (Fig. S4A-B). Taken together, this complements the RT-qPCR time courses of NGN2/ASCL1-induced neurogenesis and BMSC differentiation and demonstrates temporal heterogeneity in the level of *H19* during the differentiation process in both the human iPS cells and the human BMSCs.

### Promoter usage, transcript structure and regional OCT4/SOX2 occupancy at the *H19* locus in human pluripotent stem cells

To determine whether the *H19* variant switch involves a change in TSS, and whether this phenomenon generalizes across additional pluripotent systems, we examined publicly available CAGE data from the FANTOM5 consortium^44,45^. CAGE maps capped 5′ transcript ends and therefore provides strand-specific measurements of transcription initiation. We compared signal across the canonical *H19* promoter/exon 1 region and the broader annotated variant 3 first-exon region. CAGE profiling in undifferentiated H9 hESCs^46^ showed variant 3-associated signal and minimal signal at the canonical TSS (Figs. 4A and S6A). In the FANTOM5 H9 embryoid body-to-melanocyte differentiation time course^46^, the day-0 embryoid-body samples, which were collected after four days of embryoid-body formation, contained both canonical and variant 3-associated signal, followed by progressive loss of variant 3-associated signal during differentiation (Figs. 4B and Fig. S6B). A FANTOM5 neural differentiation time course of C11 iPSCs^46^ similarly demonstrated loss of variant 3-associated initiation and emergence of canonical initiation (Figs. 4C and Fig. S6C). Across the three FANTOM5 sample sets, 98.4% of summed variant 3-region CAGE signal localized within a 81-bp interval at hg19 chr11:2022491-2022571. Signal was distributed across multiple positions, supporting a broad initiation region rather than a single fixed TSS. Because CAGE maps transcript 5′ ends, these data identify a variant 3-associated initiation region rather than the complete structure of the initiated transcripts. Thus, FANTOM5 CAGE data independently support a pluripotency-associated alternative *H19* initiation region and generalize the variant-switch pattern across independent pluripotent differentiation systems.

**Figure 4.**
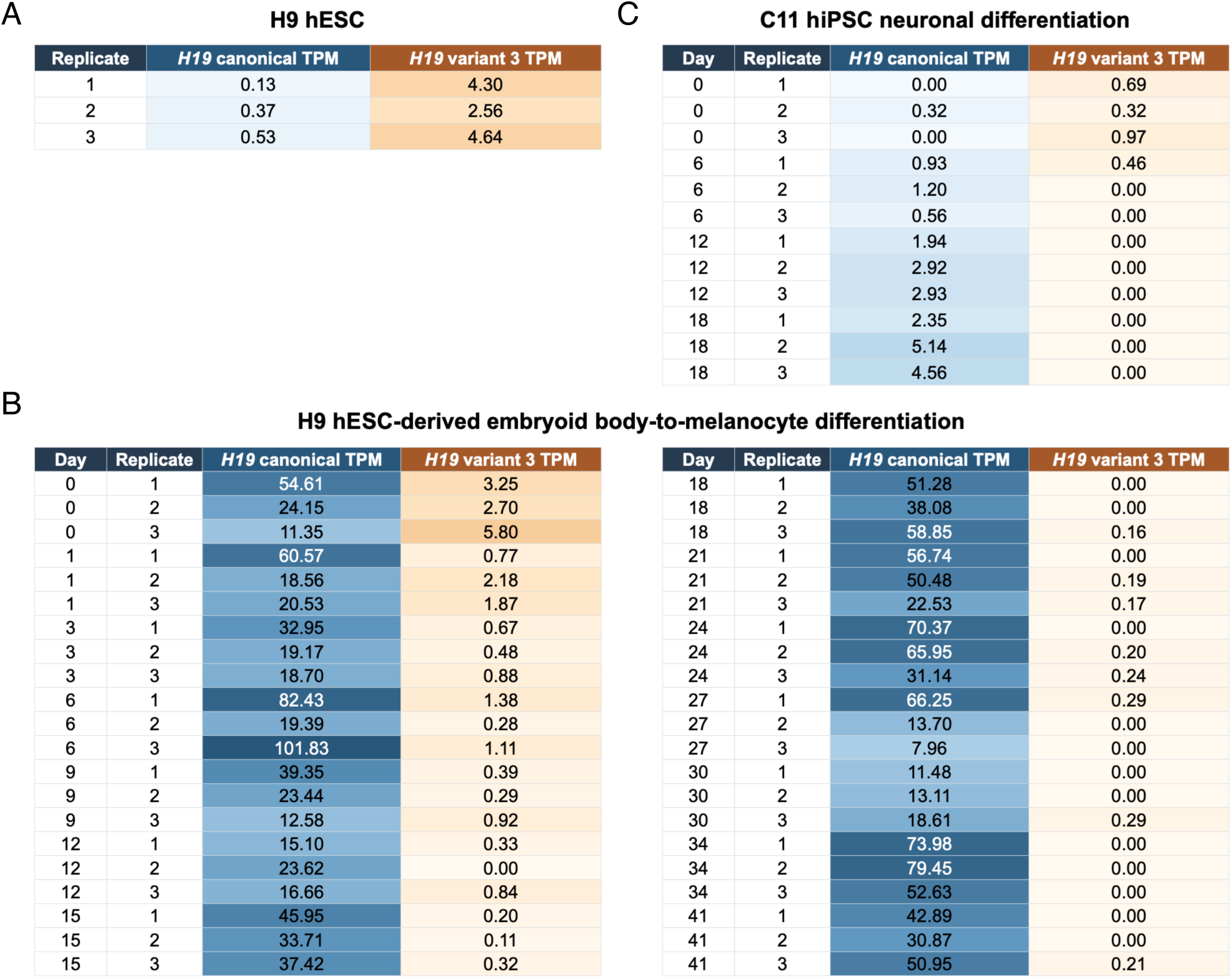
FANTOM5 CAGE supports variant 3-associated *H19* initiation in pluripotent cells and changes during differentiation. Regional reverse-strand CAGE tags per million (TPM) values are shown for **(A)** H9 hESCs, **(B)** H9-derived embryoid body-to-melanocyte differentiation, and **(C)** C11 hiPSC-to-neuron differentiation (three deposited samples per condition or time point). Canonical-associated and variant 3-associated regions correspond to the first exons of RefSeq NR_002196.3 and historical NR_131224.1, respectively. Cell shading uses a shared log₂(TPM+1) scale (blue, canonical; orange, variant 3-associated signal).

To examine whether the first exon of variant 3 was connected to all four downstream exons in pluripotent stem cells, we analyzed public long-read RNA sequencing data from KOLF2.1J iPSCs^47^ and H9 hESCs^48^. Among primary alignments with MAPQ 20-254, 21 KOLF2.1J reads and 21 H9 reads (15 in replicate 1 and six in replicate 2) contained all four expected V3 splice junctions within a 5-nt tolerance at each intron boundary (Fig. 5A). Four KOLF2.1J reads contained all four expected canonical junctions under the same filtering conditions but covered only 17–96 nt of the annotated 1,325-nt canonical first exon. No canonical-like H9 reads met these criteria (Fig. 5A). Most aligned 5′ endpoints lay near the current V3 boundary in KOLF2.1J, whereas most H9 alignments were further extended to 441-526 nt upstream of the splice junction (Fig. 5B). These findings support connection of the V3-associated first exon to the four downstream *H19* exons in both pluripotent cell lines.

**Figure 5.**
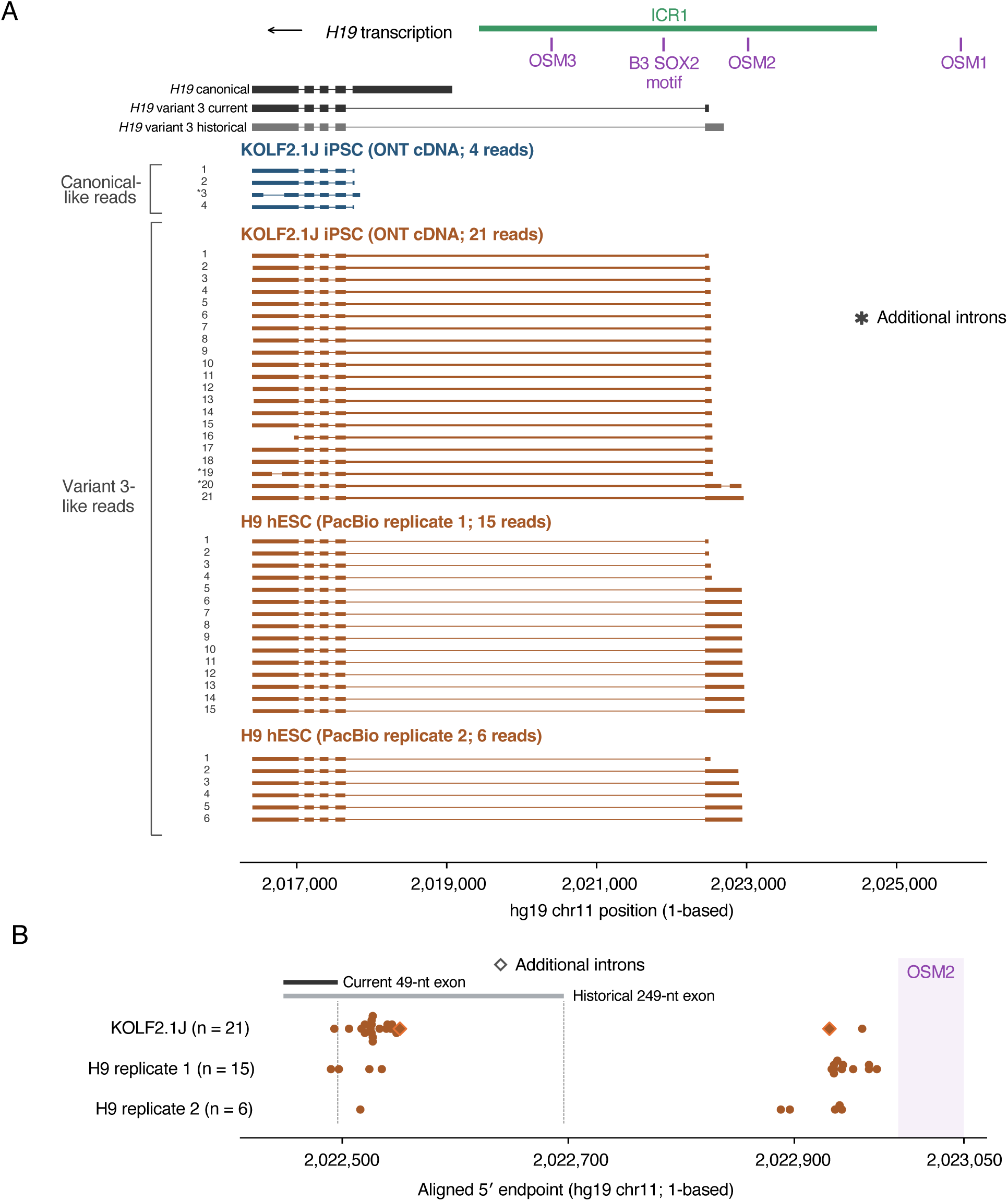
Long-read RNA-seq supports spliced *H19* variant 3-like transcripts with distinct aligned 5′ endpoints. **(A)** Canonical *H19*, current *H19* variant 3, and historical *H19* variant 3 RefSeq models above individual KOLF2.1J iPSC and H9 hESC read alignments. Displayed reads contained all four expected splice junctions for canonical *H19* or variant 3. KOLF2.1J contains four canonical-like and 21 variant 3-like reads; H9 contains 21 variant 3-like reads across two replicates (15 and six reads) and no canonical-like reads passing the alignment filters. Asterisks mark reads containing additional introns. **(B)** Aligned 5′ endpoints of variant 3-like reads relative to the current 49-nt and historical 249-nt first-exon annotations; diamonds mark reads containing additional introns.

Because loss of variant 3 coincided with loss of pluripotency factors, we next asked whether OCT4 or SOX2 occupied candidate elements in and around the human ICR1. These included three OCT4/SOX2 motifs (OSM1-3) and a separate putative SOX2 motif within the B3 repeat^42,43,49^ (Fig. 6A). Human genetic studies have identified mutations or deletions affecting these elements; notably, an A2/OSM2-region mutation was shown by gel-shift assay to impair binding of human OCT4/SOX2 proteins^42^, while complementary mouse studies demonstrated cooperative OCT4/SOX2 binding to Sox-Oct motifs and a role for these elements in maintaining ICR hypomethylation^17^. We therefore asked whether public human pluripotent-cell ChIP-seq data sets supported occupancy at these candidate regions. Processed GRCh37/hg19 ChIP-Atlas signal tracks and standardized MACS2 peak calls were examined in the UCSC Genome Browser (Fig. 6A). In the H9 hESC-to-neural progenitor study^50^, the hESC SOX2 library contained a formal peak overlapping OSM1, whereas the H9-derived neural progenitor comparator did not (Fig. 6A). In PGP1 iPSCs profiled by ENCODE^51^, one of two OCT4 libraries contained an OSM1-overlapping peak, whereas no corresponding peak call was present in the day-4 NGN1/NGN2-derived neuronal comparator (Fig. 6A). In HUES64 hESCs from the hESC-to-ectoderm study^52^, one of three OCT4 libraries contained an OSM1-overlapping peak, whereas the day-5 ectoderm comparator did not (Fig. 6A). Across these representative series of different pluripotent cells transitioning to a differentiated state, OSM1 peak calls were present in the pluripotent samples but were absent from the differentiated comparator tracks. We therefore expanded the analysis to 18 OCT4 and 14 SOX2 undifferentiated hESC/iPSC studies comprising 69 biological-sample groups. At the study level, OSM1 peak prevalence was 50.0% for OCT4 and 35.7% for SOX2, whereas OSM2 prevalence was 5.6% and 21.4%, respectively (Fig. 6B). No study-level peaks overlapped OSM3 or the B3 SOX2 motif for either factor (Fig. 6B). Biological-sample-level analysis produced the same overall site-preference pattern suggesting that the OSM1 site is the predominant site for binding in pluripotent cells with less-frequent binding at OSM2 (Fig. S7A).

**Figure 6.**
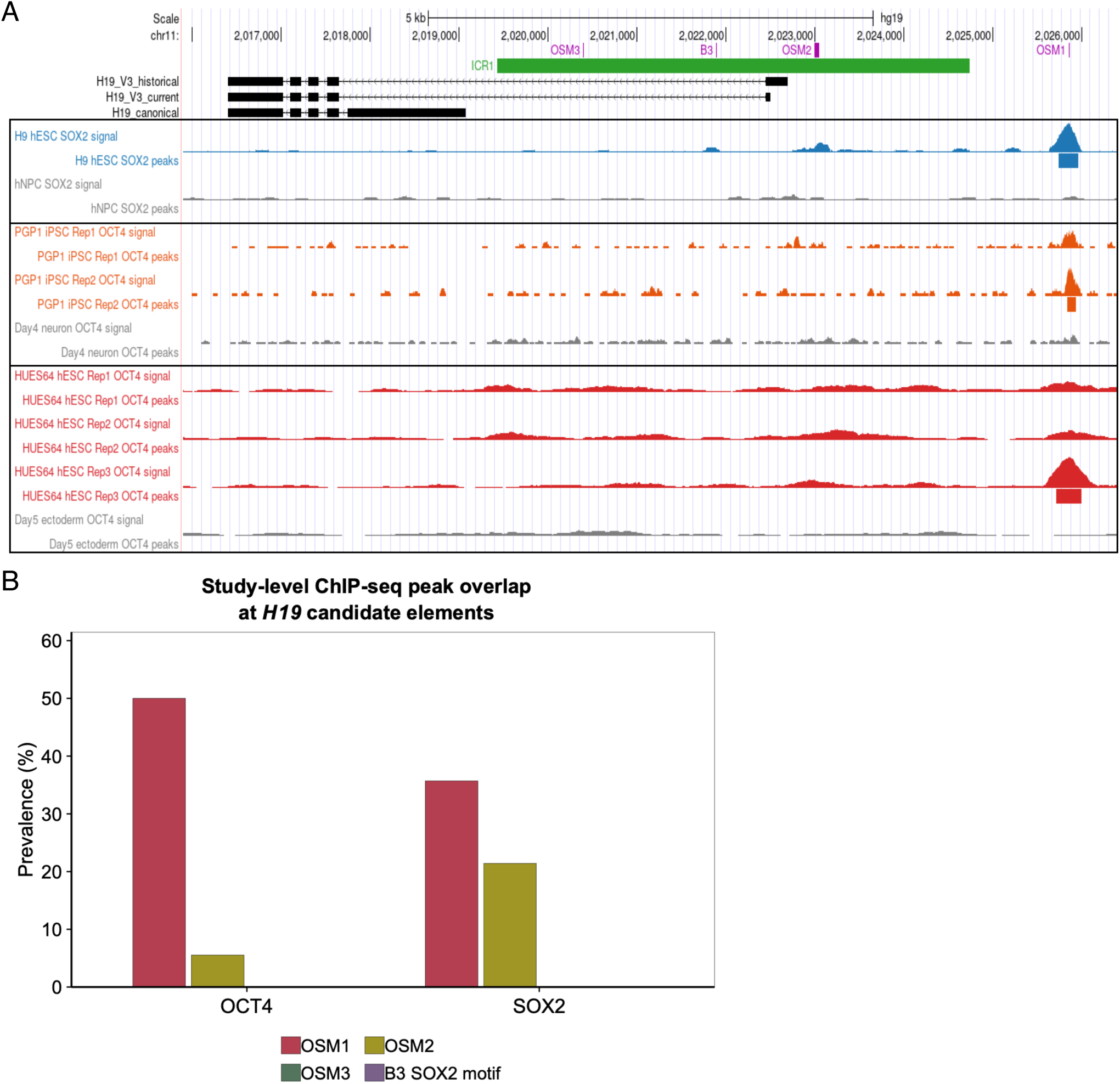
Public OCT4 and SOX2 ChIP-seq peak overlap at candidate *H19* regulatory elements. **(A)** Representative ChIP-Atlas signal tracks and MACS2 peak calls (q<1×10⁻⁵) across the *H19/IGF2* locus, including ICR1 and the OSM1–3 and B3 SOX2 candidate motifs, in pluripotent and differentiated comparator samples. **(B)** Study-level prevalence of peaks overlapping each candidate element in eligible undifferentiated hESC/hiPSC studies (18 OCT4 studies and 14 SOX2 studies). A study was positive if at least one eligible biological sample contained an overlapping peak.

Expanded-window sensitivity analyses using uniform 101-bp and 201-bp windows centered on each candidate element produced essentially the same site-prevalence pattern as the direct-overlap analysis (Fig. S7B-C). One additional OCT4 experiment contained a significant ChIP-Atlas MACS2 peak within both expanded B3-centered windows; this peak was near, but did not directly overlap, the literature-defined B3 motif. No other site calls changed. Overall, these data support a cross-study convergence of pluripotency-factor ChIP-seq occupancy at *H19* candidate regions, with the strongest recurrent evidence for OCT4 at OSM1, recurrent SOX2 occupancy at OSM1 and OSM2, and little evidence for consistent occupancy of either factor at OSM3 or the B3 SOX2 motif in human pluripotent cells.

## Discussion

In this study we identify a novel shift in *H19* isoform usage during differentiation of human pluripotent stem cells. Variant 3, an understudied isoform, was the predominant *H19* transcript in both hiPSC and hESC lines. Levels of variant 3 declined rapidly, falling below LLRQ upon loss of pluripotency during NGN2/ASCL1-induced neurogenesis. Simultaneously, the canonical *H19* transcript and *IGF2* transcripts increased, although the increase was transient. In contrast, variant 3 was below LLRQ in multipotent BMSCs, although the canonical *H19* transcript was detected at baseline and was markedly increased during induced neurogenic and osteogenic differentiation. Similarly, variant 3 was below LLRQ in adult primary and cancer-derived cell types; thus, variant 3 dominance was associated with pluripotency in the cell types examined. Therefore, *H19* regulation involves not only changes in total *H19* levels, but also cell-state-dependent changes in dominant detected isoform usage.

Canonical *H19* and *IGF2* showed different expression responses across different differentiation conditions. In our KOLF2.1J iNeuron model we observed a coordinated, transient induction of both canonical *H19* and *IGF2* during early neurogenesis (Fig. 2D). The BMSC neurogenic induction produced a pronounced increase in *H19* levels, while adipogenic induction produced a pronounced increase in *IGF2* transcripts. The BMSC osteogenic induction resulted in an increase in both *H19* and *IGF2* (Fig. 3A).

Additionally, while *H19* and *IGF2* rose and fell during neurogenesis in the KOLF2.1J iNeuron model by day 14 these transcripts remained elevated during BMSC differentiation at day 14. The difference between forced upregulation of transcription factors in the iNeurons and media-based induction of BMSCs may explain the temporal dynamics of *H19/IGF2* transcripts and may represent differences in the progression of differentiation as the inducible system rapidly elevates neural specific factors which are likely to increase in a more prolonged timeframe during directed differentiation using exogenous factors. Given the variations in levels in the different differentiation protocols, the magnitude and timing of *H19* and *IGF2* induction appears to be dependent on differentiation context, and these transcripts are not uniformly coupled.

RNAscope complements the bulk RT-qPCR data by visualizing variation in *H19* signal between individual cells. Undifferentiated KOLF2.1J cells contained uniformly low signal across all cells, while day 5 iNeuron cultures contain a mix of high, moderate, and low *H19* containing cells (Fig. S4A). This may represent temporal differences in *H19* response to NGN2/ASCL1-induced neurogenesis and/or the extent to which *H19* responds to NGN2/ASCL1 across individual cells. BMSCs, however, demonstrate heterogeneity at baseline with a small subpopulation of cells containing moderate-to-high *H19* signal while the majority show low signal (Fig. S4B-C). Upon neurogenic and osteogenic differentiation, *H19* levels increased dramatically but was heterogeneous across cells (Fig. S4B). This heterogeneity may indicate a requirement for *H19* at a specific transition point during the differentiation process although further study is required to fully understand the implications of this heterogeneity.

The public FANTOM5 datasets independently reinforced the RT-qPCR results, as undifferentiated hESC and iPSC samples predominantly showed variant 3-associated signal. While the H9 hESC cells contain prominently variant 3-associated signal, the canonical *H19*-associated signal is already prominent within the four-day H9-derived embryoid bodies and before subsequent melanocyte differentiation, suggesting that this shift occurs early during differentiation (Fig. 4A-B). The variant 3-associated signal progressively declines upon melanocyte differentiation (Fig. 4B). Additionally, the C11 iPSC-to-neuron differentiation shows declining variant 3-associated signal and increasing canonical *H19*-associated signal (Fig. 4C). Together, this data further supports the notion that this phenomenon occurs in hESC and iPSC samples and extends beyond the KOLF2.1J iNeuron model. This data is further supported by the long-read RNA-seq, which shows that most reads meeting the quality and complete V1/V3 junction analysis criteria were variant 3-like molecules in KOLF2.1J iPSC and H9 hESC samples (Fig. 5A). Interestingly, we observed that while the *H19* transcripts from KOLF2.1J cells primarily contained a short exon 1 similar to the current variant 3 annotation (NR_131224.2), the majority of transcripts in H9 cells had aligned exon 1 reads spanning 441-526 nt (Fig. 5B). CAGE signal farther upstream was also visible in all three datasets (Fig. S6A-C) but fell outside the annotated variant 3 exon 1 interval used for quantification as we had not established whether transcripts initiating from that region connected to downstream exons during the initial analysis.

The independent OCT4 and SOX2 ChIP-seq analysis further identified OSM1 as the candidate region with the strongest recurrent peak overlap in pluripotent cells (Fig. 6A-B); corresponding peak calls were absent in the differentiated comparator datasets examined (Fig. 6A). Peaks for both transcription factors also overlapped OSM2, although less frequently than OSM1 (Fig. 6B). OSM1 lies approximately 3.4 kb upstream of the current annotated variant 3 initiation region, while OSM2 lies approximately 500 bp upstream of that region (Fig. 6A). OSM2 is therefore in close proximity to the upstream CAGE signal and extended aligned 5’ endpoints of variant 3-like reads (Figs. S6A-C and 5B). Additionally, the decline in variant 3 levels coincided temporally with loss of *POU5F1* and *SOX2* transcript levels during NGN2/ASCL1-induced neurogenesis in KOLF2.1J cells (Figs. 1C and 2D). Interestingly, beyond their role in maintaining ICR hypomethylation, murine ICR Sox-Oct motifs have been shown to stimulate *H19* promoter activity in luciferase reporter assays in mouse embryonic carcinoma cells^17^. This transcriptional activation was abolished by mutation of either the OCT-or SOX-binding sites within the composite motifs and markedly reduced following OCT4 knockdown. Together, these findings support the hypothesis that changes in pluripotency-associated transcription factor occupancy, potentially accompanied by changes in chromatin organization at the *H19* locus, contribute to the shift in *H19* isoform usage during early differentiation. Additional mechanisms may further govern the upregulation of canonical *H19* that we observed across various differentiation lineages, as upregulation of canonical *H19* also occurred during BMSC differentiation. We speculate that occupancy at OSM1 and/or OSM2 may contribute to variant 3 initiation-site usage.

Particularly notable is the substantial sequence difference between variant 3 and the canonical-like transcripts. We hypothesize that NCBI variants 1, 2, 4, 5, 6, and 7 can arise from alternative splicing of transcripts initiated at the canonical promoter whereas alternative TSS usage for variant 3 is supported by our CAGE analysis. A few previous studies have identified *H19* variant 3 in non-pluripotent contexts, including glioma and mesothelial cells^53,54^; however, expression was minimal in comparison to canonical-like variants^54^. Interestingly, a comparative isoform analysis showed that the variant 3-like exon 1 annotation is shared between primates although a cognate in mice is not annotated^55^. Thus, the reported difference warrants caution in extrapolating isoform-specific findings between species.

The functional consequences of choosing variant 3 rather than canonical *H19* remain unknown. One certainty about variant 3 is that due to the lack of canonical exon 1 it cannot serve as a precursor for miR-675^22^; thus, exclusion of canonical exon 1 may be a functionally important distinction that could be shared between variant 3-like transcripts with either longer or shorter first exons. In a similar respect, shared downstream exons may support common functions between variant 3-like molecules with different length first exons during pluripotency. Additionally, different length first exons of variant 3 could be functionally redundant due to structural interactions of the 3’ end with downstream exons. The previously described GMRSP peptide that has been shown to be produced from an ORF specific to variant 3 raises the possibility that variant 3 may also function entirely or in-part by producing a protein in pluripotent cells^37^. Prior work supports both nuclear and cytoplasmic activities of *H19*, including recruitment of MBD1 to chromatin^56^, interaction with RBFOX2/hnRNPM in splicing regulation^57^, and antagonism of let-7 microRNAs^58^. Interestingly, previous work has indicated that knockdown of *H19* in pluripotent cells has negative effects on pluripotency, although experimentally the study targeted shared exon 5 and thus could not distinguish isoform-specific effects^31^. Although not specifically related to *H19* transcripts, it has been shown that low-level transcription through the ICR1 on the paternal allele is necessary for establishment of methylation in mouse prospermatogonia^59^. Together, these findings raise three non-exclusive possibilities: variant 3 may retain or acquire mature-RNA functions through its shared and unique sequence and structure; it may produce GMRSP in pluripotent cells; and/or transcription from the variant 3 region may influence regulation of the *H19/IGF2* locus independently of the mature transcript.

The SYBR Green assay incorporates assay-specific RNA calibration and efficiency-adjusted *H19* quantification in an attempt to best adhere to the Minimum Information for Publication of Quantitative Real-Time PCR Experiments (MIQE)^60^. We validated the canonical and variant 3 assays across a wide range of copies/reaction input which showed linear relationships for both, with no expected specific melt peaks against the noncognate template, nor amplification within no-template controls. The assay-specific standard curves were used to convert C_T_ values to estimated copies per reaction. While we were carefully conservative about defining LLRQ in the assay to only reactions with dominant specific melt peaks even a single peak does not exclude a fluorescence contribution from nonspecific products. As well, copy estimates assume an average of 10 pg total RNA per cell. The use of a single iPSC line for the neurogenesis to demonstrate the variant 3-to-canonical switch is a limitation although CAGE analysis further generalized this phenomenon; long-read RNA-seq additionally supported variant 3 dominance in KOLF2.1J and H9 cells. Causal regulation by OCT4/SOX2 and functional requirements of variant 3 remain untested.

The present study establishes a cell-state-dependent change in dominant *H19* isoform usage but does not determine whether variant 3 itself contributes to pluripotency or whether the isoform transition is functionally required for differentiation. Future work requires isoform-specific functional studies to distinguish precise roles of canonical *H19* and variant 3. Overall, these findings establish *H19* isoform selection as an additional layer of regulatory complexity at the *H19/IGF2* locus during cell-state transitions.

## Materials and Methods

### Cell culture

All cells grown in our laboratory were cultured at 37 °C and 5% CO2. Parental KOLF2.1J human induced pluripotent stem cells (hiPSCs) were purchased from The Jackson Laboratory (JIPSC001000). Parental KOLF2.1J and KOLF2.1J-C8 hiPSCs were maintained in StemFlex Medium (Gibco, A3349401) on plates coated with Synthemax II-SC Substrate (Corning, 3535). The medium was changed every two days. Cells were passaged as clumps using ReLeSR (STEMCELL Technologies, 05872) at approximately 80% confluence, typically at a 1:6 split ratio.

Human bone marrow-derived mesenchymal stem cells (BMSCs) were purchased from ScienCell (7500) and maintained on uncoated tissue culture-treated plates in Mesenchymal Stem Cell Growth Medium 2 (PromoCell, C-28009). Cells were detached with Accutase (Sigma-Aldrich, A6964) at approximately 80% confluence and reseeded at 4 × 103 cells/cm2.

U251 and LNCaP cells were provided by Dr. Srinivas Somarowthu and Dr. Alessandro Fatatis, respectively. Both lines were cultured in Eagle’s minimum essential medium with Earle’s salts and L-glutamine (Corning, 10-010-CV). Each bottle was supplemented with 50 mL of GemCell fetal bovine serum (FBS, GeminiBio, 100-500), 10 mL of 50× MEM Amino Acids without L-glutamine (Corning, 25-030-CI), and 5 mL each of 100× MEM Vitamin Solution (Gibco, 11120-052), 100 mM sodium pyruvate (Gibco, 11360-070), and 200 mM L-glutamine (Corning, 25-005-CI). Lenti-X 293T cells (Takara Bio, 632180) were cultured in HyClone DMEM/F-12 (1:1, Cytiva, SH30126.01) supplemented with 50 mL FBS and 5 mL of 200 mM L-glutamine per bottle.

### Generation of the doxycycline-inducible NGN2/ASCL1 iPSC line

Parental KOLF2.1J hiPSCs were transduced with the doxycycline-inducible NGN2/ASCL1 construct pLVX-UbC-rtTA-Ngn2:2A:Ascl1 that was created in the lab of Dr. Fred Gage (UNA, Addgene plasmid 127289)^40^. Lentiviral particles were generated in Lenti-X 293T cells using the PcPack2 packaging system. Following puromycin selection, 12 colonies were picked individually and eight expanded sufficiently for screening. Expanded clones were screened for seven days after doxycycline induction. KOLF2.1J-C8 was selected based on neuronal morphology and uniformity of differentiation.

### NGN2/ASCL1-induced neuronal differentiation

On day 0, doxycycline was added to the existing StemFlex Medium in KOLF2.1J-C8 cultures to a final concentration of 10 µg/mL. On day 1, the medium was replaced with fresh StemFlex containing 10 µg/mL doxycycline. On day 2, cells were switched to a 1:1 mixture of Neurobasal Medium (Gibco) and StemFlex Medium supplemented with 1× N2 Supplement-A (STEMCELL Technologies, 07152), 1× B-27 Supplement (Gibco, 17504044), 0.5 µg/mL iMatrix-511 silk (Nippi, 892-021), 10 ng/mL recombinant human/mouse/rat BDNF (PeproTech, 450-02-100UG), and 10 µg/mL doxycycline.

On day 3, cells were dissociated with Accutase and seeded at 3 × 104 cells/cm2 on plates coated with poly-D-lysine (Gibco, A3890401) and iMatrix-511 silk. The week 1 maturation medium consisted of Neurobasal Medium supplemented with 1× B-27, 0.5 µg/mL iMatrix-511 silk, 10 ng/mL BDNF, 10 ng/mL recombinant human NT-3 (RayBiotech, 230-00566), 10 µg/mL doxycycline, and 10 µM 5-ethynyl-2′-deoxyuridine (EdU, Invitrogen, A10044). RevitaCell Supplement (Gibco) was added at seeding. Cultures were fed with fresh week 1 maturation medium on day 6. Beginning on day 10, the fresh maturation medium contained 2 µg/mL doxycycline and 1 µM EdU, with the other maturation supplements at the same concentrations. At each feed after seeding, we removed about half the remaining medium and added fresh medium to restore the original volume. Cells were analyzed through day 14. Phase-contrast images were acquired on days 0, 5, and 14 with a ZEISS Primovert inverted microscope and AxioCam ERc 5s camera.

### BMSC differentiation

BMSCs were seeded on plates coated with human fibronectin (1 µg/cm2, PromoCell, C-43060) for neurogenic and adipogenic differentiation or rat-tail collagen I (Enzo Life Sciences, ALX-522-440-0050) for osteogenic differentiation. Undifferentiated BMSCs were collected on day 0. Cells were cultured for 14 days in the corresponding PromoCell differentiation medium (neurogenic, C-28015, adipogenic, C-28016, osteogenic, C-28013), with medium changes twice weekly.

### RNA sources and extraction

Parental KOLF2.1J cells, KOLF2.1J-C8 cells and their neuronal derivatives, BMSCs, Lenti-X 293T, U251, and LNCaP cells were lysed in TRIzol (Invitrogen, 15596026). Total RNA was isolated using the Direct-zol RNA Miniprep Kit (Zymo Research, R2052) according to the manufacturer’s protocol, including on-column DNase I treatment. RNA was eluted in 50 µL of water and quantified with the Qubit RNA Broad Range assay (Thermo Fisher Scientific).

HuES8 RNA was provided by Ridhdhi Desai. RNA from human cardiac fibroblasts (HCFs), astrocytes, and senescent astrocytes had been collected previously in our laboratory; the astrocyte RNA was collected as part of a previous study^61^. HCFs and astrocytes were purchased from ScienCell (6300 and 1800, respectively).

### RT-qPCR

One-step reverse transcription quantitative PCR (RT-qPCR) was performed on an Applied Biosystems 7500 Real-Time PCR System. Reactions containing cellular RNA used 20 ng of total RNA per well. For assays normalized to 18S rRNA, we measured 18S in separate wells on the same plate with a 1× TaqMan assay (Thermo Fisher Scientific, Hs03928989_g1) using the same RNA input, master mix, volume, and cycling conditions as the targets.

Plates containing SYBR Green assays were run in 25 µL reactions using the Verso SYBR Green 1-Step qRT-PCR Low ROX Kit (Thermo Scientific, AB4106C). SYBR Green reactions contained 100 nM of each primer (Fig. S1B). Reverse transcription at 50 °C for 15 min was followed by polymerase activation at 95 °C for 15 min and 40 cycles of 95 °C for 15 s, 60 °C for 30 s, and 72 °C for 30 s. Fluorescence was collected at the end of each extension step. For melt curve analysis, reactions were held at 95 °C for 30 s and 60 °C for 1 min. Fluorescence was then collected continuously as the temperature increased from 60 to 95 °C at the instrument’s 1% ramp setting.

Plates containing only TaqMan assays were run in 20 µL reactions using KiCqStart One-Step Probe RT-qPCR ReadyMix, Low ROX (Sigma-Aldrich, KCQS08), with each assay at 1×. Reverse transcription at 50 °C for 1 min was followed by initial denaturation at 95 °C for 1 min and 40 cycles of 95 °C for 10 s and 60 °C for 60 s. Fluorescence was collected at the end of each 60 °C step. The target assays (Thermo Fisher Scientific) were *POU5F1* (Hs04260367_gH), *SOX2* (Hs04234836_s1), *NANOG* (Hs02387400_g1), and *MAP2* (Hs00258900_m1).

The preliminary primer screen assessed amplification, melt profiles, and product size without reference-gene normalization. The *H19* variant 3 amplicon of the expected size from day-0 KOLF2.1J-C8 cells was excised from the gel, purified using the GenElute Gel Extraction Kit (Sigma-Aldrich, NA1111-1KT), reamplified with the variant 3 primers, and sequenced by Plasmidsaurus. The sequence was compared with *H19* reference sequences using BLAST. For expression calls, a specific melt peak alone or dominant over nonspecific peaks indicated detection. Samples with an absent or nondominant specific peak were classified as below the lower limit of reliable quantification (LLRQ).

### *H19* RNA standards and standard curves

Full-length human *H19* variant 1 (RefSeq NR_002196.2) was amplified from plasmid DNA using Platinum II Hot-Start PCR Master Mix (Thermo Fisher Scientific) and the following primers (5′ to 3′): forward, ACGTGAATTCTAATACGACTCACTATAGGGAGTTAGAAAAAGCCCGGGCTAG, and reverse, TACGGGATCCGCTGTAACAGTGTTTATTGATG. The forward primer introduced a T7 promoter upstream of the insert. Full-length *H19* variant 3 (RefSeq NR_131224.2) was obtained as a synthetic gBlocks DNA fragment (Integrated DNA Technologies) containing an upstream T7 promoter and flanking restriction sites. Both inserts were cloned into a plasmid backbone provided by Srinivas Somarowthu using EcoRI-HF and BamHI-HF (New England Biolabs). Colonies were screened by colony PCR, and plasmids from selected colonies were sequenced by Plasmidsaurus.

The plasmids were linearized with BamHI downstream of the *H19* inserts and transcribed using the TranscriptAid T7 High Yield Transcription Kit (Thermo Fisher Scientific). After transcription, the reactions were treated with RNase-free DNase I to remove template DNA. RNA was purified using the RNA Clean & Concentrator-25 kit (Zymo Research). Purified RNA was quantified with the Qubit RNA Broad Range assay and assessed on a 1% agarose gel after denaturation in formamide loading buffer.

Copy concentrations were estimated from RNA mass concentrations using lengths of 2,315 nt for canonical *H19* and 1,039 nt for variant 3 and an assumed average molar mass of 340 g/mol per nucleotide. Each stock was diluted in water to prepare a five-point, tenfold dilution series with nominal inputs of 103 to 107 copies per reaction. Each dilution was run in technical duplicate under the SYBR Green cycling and melt curve conditions described above. Each primer pair was also tested against the opposite *H19* variant to assess cross-reactivity. No-template controls contained water. Standard curves were fitted by linear regression of mean C_T_ against log10 of nominal copies per reaction. Amplification efficiency (%) was calculated as 100 × [10−1/slope − 1].

### Immunofluorescence

Day 10 iNeurons were fixed in 4% paraformaldehyde for 10 min, permeabilized with 0.1% Triton X-100 for 10 min, and blocked for 1 h in PBS containing 1% bovine serum albumin, 0.30 M glycine, and 0.1% Tween 20. All incubations were at room temperature.

Separate wells were incubated for 1 h with either mouse monoclonal anti-MAP2 (clone M13, Invitrogen, 13-1500, 1:400) or mouse monoclonal anti-beta III tubulin (clone 2G10, Sigma-Aldrich, T8578, 1:1000) in blocking buffer. After washing with PBS, cells were incubated for 1 h in the dark with Alexa Fluor 488-conjugated rabbit anti-mouse IgG (H+L) (Invitrogen, A-11059, 1:2000) in blocking buffer. Cells were washed with PBS and counterstained with 1 µg/mL DAPI for 1 min. Fluorescence images were acquired on an EVOS fluorescence microscope (Thermo Fisher Scientific) using a 20× objective.

### RNAscope in situ hybridization

RNAscope was performed on KOLF2.1J-C8 cultures at days 0, 5, and 14 of neuronal differentiation, undifferentiated day-0 BMSCs, and day-14 neurogenic and osteogenic BMSCs. Probe controls used KOLF2.1J-C8 cells. Cells were cultured on 8-well Lab-Tek glass chamber slides (Nunc, 177402) and prepared according to the Advanced Cell Diagnostics RNAscope protocol for cultured adherent cells, with fixation in 4% paraformaldehyde for 30 min at room temperature. Protease III (Advanced Cell Diagnostics, 322381) was used at 1:15 for all cell types.

Single-channel detection was performed using RNAscope Multiplex Fluorescent Detection Reagents v2 (Advanced Cell Diagnostics, 323110) according to the manufacturer’s instructions. We used the Hs-H19 probe (C1, 400771) to detect *H19* RNA across isoforms. Hs-POLR2A-C3 (310451-C3) served as the positive control. The RNAscope 3-plex dapB probe (320871) served as the negative control. The H19 and dapB probes were developed with HRP-C1 and the POLR2A probe with HRP-C3. Opal 570 (Akoya Biosciences, OP-001003) was diluted 1:750 in RNAscope TSA buffer. Slides were counterstained with DAPI and mounted with ProLong Gold Antifade Mountant.

Slides were imaged 8 h to 2 weeks after mounting on an Olympus BX61 fluorescence microscope equipped with a Hamamatsu C4742-80-12AG camera. Representative images were acquired using 20×, 40×, and 60× objectives and assessed qualitatively. Display brightness was increased by 50% in PowerPoint for the panels specified in the legends to Figs. S4 and S5.

### RT-qPCR data analysis and statistics

RT-qPCR data were analyzed in R (Supplementary Data 1). For the differentiation time courses, expression was normalized to 18S rRNA and reported relative to day 0. Fold changes for canonical *H19* and variant 3 were calculated using the amplification efficiency measured for each assay. We assumed 100% efficiency for *IGF2* and 18S rRNA.

For KOLF2.1J-C8 neurogenesis, three biological replicates per time point were run in technical duplicate. Duplicate C_T_ values were averaged before analysis. Each later time point was compared with day 0 by one-way ANOVA with two-sided Dunnett comparisons (emmeans, multivariate t adjustment). For *POU5F1*, *SOX2*, *NANOG*, and *MAP2*, the comparisons used ΔC_T_. For the *H19* isoforms and *IGF2*, they used log2 fold changes calculated with the assay efficiencies described above. We report geometric means and standard deviations across biological replicates and considered adjusted P < 0.05 significant.

For BMSC differentiation, one biological replicate per lineage was run in technical triplicate. Fold changes were calculated for measurable wells using the sample’s mean reference C_T_ and summarized as the mean and standard deviation across technical wells without significance testing.

For each biological replicate in the KOLF2.1J-C8 time course, we estimated canonical *H19* and variant 3 copies per reaction from the mean target C_T_ using standard curves fitted to individual standard wells. We divided by 2,000 to estimate copies per cell equivalent (20 ng RNA per reaction, assuming 10 pg RNA per cell). Time-point means are marked [E] if any replicate fell outside the calibration range. Estimates below the LLRQ were retained as exploratory. For the day 0, 5, and 14 pie charts, we expressed each isoform as a percentage of the two isoforms’ combined estimated copies in each biological sample, then averaged the percentages across biological replicates.

### FANTOM5 CAGE analysis

Processed GRCh37/hg19 FANTOM5 CAGE tracks reported in tags per million (TPM)^44–46,62^ were analyzed for H9 hESCs, H9 embryoid body-to-melanocyte differentiation, and C11 iPSC-to-neuron differentiation. Reverse-strand tracks were analyzed separately for each deposited sample. Quantification regions were defined from current and historical *H19* RefSeq annotations^21^. Canonical *H19* was quantified across its first exon; variant 3 was quantified across its historical first-exon interval, with separate totals for the current exon and historical extension.

Signal was extracted using UCSC bigWigToBedGraph^63^. Regional TPM was calculated by summing track values weighted by the number of overlapping bases, without additional filtering or normalization. Nonzero variant 3 CTSS ranges and pooled signal fractions within the predefined dominant interval were summarized. Table shading used log₂(TPM + 1). Sample results, interval definitions, CTSS contributions, and sources are provided in Supplementary Data 2.

### Long-read RNA sequencing analysis

Public KOLF2.1J iPSC ONT cDNA alignments^47^ and H9 hESC PacBio cDNA alignments^48^ were queried by index using samtools v1.21. Extraction used the deposited native references, including the H9 haplotype-1 assembly^64^.

Canonical *H19* and variant 3 RefSeq transcripts^21^ were aligned to hg19 and the native references using minimap2 v2.31-r1302 (splice:hq)^65^ to define junction models. Primary alignments with MAPQ 20–254 were retained when all four expected junctions of either model were present, with donor and acceptor boundaries each within 5 nt of the model. Unmapped, secondary, and supplementary alignments were excluded. Full first-exon coverage was not required; additional introns were retained and marked, and exact junction matches were recorded separately.

Native genomic windows were aligned to hg19 using minimap2 (asm5)^65,66^ to transfer junction coordinates and read exon blocks between references. Selection retained the original RNA mapping qualities. For negative-strand *H19*, the aligned 5′ endpoint was the highest-coordinate aligned base and does not establish a capped transcription start site. Read-level results, junction models, coordinate conventions, reference accessions, and sources are provided in Supplementary Data 3.

### OCT4 and SOX2 ChIP-seq analysis

Processed GRCh37/hg19 OCT4 and SOX2 ChIP-seq data were obtained from ChIP-Atlas^67,68^. The prevalence cohort comprised undifferentiated hESC/iPSC samples, including defined pluripotency states, with standardized MACS2 peak calls at q < 1 × 10⁻⁵ (threshold 05). Differentiated derivatives, reprogramming intermediates, explicitly perturbed samples, artificial mixtures, and records lacking these peak calls were excluded. Retained studies^50,52,69–92^ and experiment eligibility are documented in Supplementary Data 4.

Peak overlap was evaluated at the literature-derived OSM1–3 and B3 SOX2 elements^17,42,43,49^. An experiment was positive when a threshold-05 peak overlapped the candidate interval by at least one base. Sensitivity analyses used 101-bp and 201-bp windows centered on each interval’s midpoint, using the lower central base for even-length intervals.

SRX021070 and SRX021071 were collapsed as technical replicates of one biological sample, positive if either experiment was positive. Other retained experiments were treated as separate biological samples. A study was positive if any eligible biological sample was positive; prevalence was calculated across eligible studies and, separately, biological samples after technical-replicate collapse.

Representative signal tracks and corresponding peaks were displayed in the UCSC Genome Browser^66^. Comparisons comprised H9 hESCs versus neural progenitors^50^, PGP1 iPSCs versus induced neurons^51^, and HUES64 hESCs versus ectoderm^52^. Differentiated comparators were excluded from prevalence calculations, and signal heights were not quantitatively compared across studies. Interval definitions, experiment-and study-level calls, prevalence summaries, displayed libraries, and sources are provided in Supplementary Data 4.

## Acknowledgements

This work is supported by NIH grants AG071815 and AG102492. ChatGPT (OpenAI) was used to revise manuscript wording and to assist in drafting and revising R scripts for data analysis and visualization. The authors take responsibility for the final manuscript and reported results.

