## Supplemental Figure S1 for "*H19* isoform switching during neuronal differentiation reveals a distinct pluripotency-associated non-canonical transcript"

A

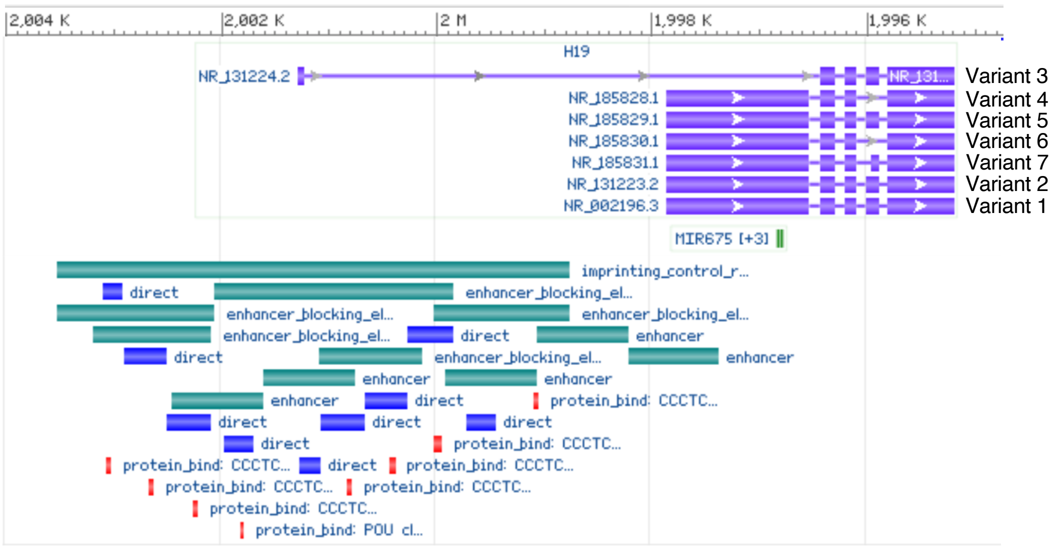


B

| Assay | Forward primer sequence (5'-3') | Forward primer location | Reverse primer sequence (5'-3') | Reverse primer location |
|---|---|---|---|---|
| All canonical-like | TTCGGGCTGGAGACGAG | Exon 1 | CCATGTCATGTCCTGCTTGT | Exon 2 |
| Variant 2 | ATCGGCTCTGGAAGCTAGA | Exon 4 | GGTGGCTGGTGGTCAAC | Exon 5 |
| ***H19* variant 3 (screen and final assay)** | CAGATGCCACGGAATCAGTT | Exon 1 | CCTGCTTGTCACGTCCAC | Exon 2 |
| Variant 4 + 6 | AGCGGGTCTGTTTCTTTACTT | Exon 3 | GGTTCCTCTAGCTTCACCTTTC | Exon 4 |
| Variant 4 + 5 | GAGACGAGGGTCTCCAG | Exon 1 | CATGTCATGTCCTGCTTGT | Exon 2 |
| Variant 7 | GTCTGTTTCTTTACTTCCTCCAC | Exon 3 | CAGCATATTCATTTCCAAGCTTTC | Exon 4 |
| ***H19* canonical (final assay)** | ATCGGTGCCTCAGCGTTC | Exon 1 | TCCTGCTTGTCACGTCCAC | Exon 2 |
| ***IGF2* (final assay)** | TTGGACAGAAGCCCAAAGAG | Exon 4 | CTGACGAATGGGCAGGTAAT | Exon 4 |

C

88°C specific product

86°C specific product

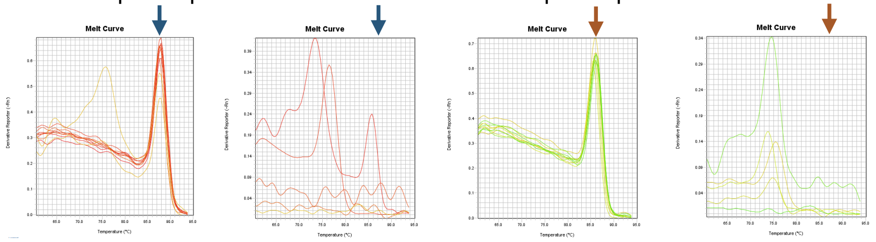


Variant 1 RNA, canonical assay

Variant 1 RNA, variant 3 assay

Variant 3 RNA, variant 3 assay

Variant 3 RNA, canonical assay

D

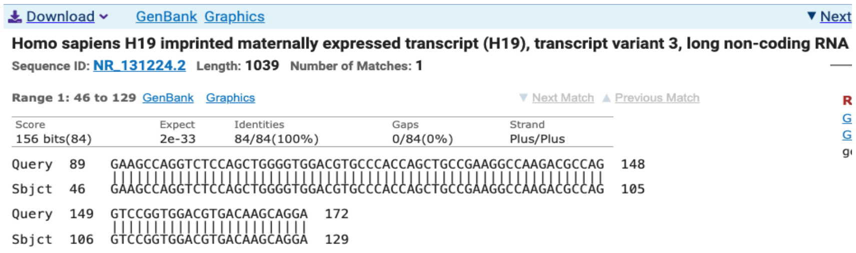


**Figure S1. Design and validation of *H19* isoform-junction SYBR Green RT-qPCR assays.** **(A)** NCBI genome-browser view of the *H19*/*IGF2* locus, annotated *H19* transcript variants, and ICR1 regulatory features. **(B)** Primer sequences and exon locations for the preliminary *H19* transcript screen, optimized canonical *H19* and variant 3 assays, and *IGF2* assay. **(C)** Melt-curve profiles of the optimized *H19* assays tested against IVT canonical *H19* and variant 3 RNA, with expected specific peaks near 88°C and 86°C, respectively. Blue and orange arrows mark canonical *H19* and variant 3-specific peaks, respectively. **(D)** NCBI BLAST alignment confirming the identity of the sequenced, gel-purified variant 3 amplicon.
