## Supplemental Figure S2 for "*H19* isoform switching during neuronal differentiation reveals a distinct pluripotency-associated non-canonical transcript"

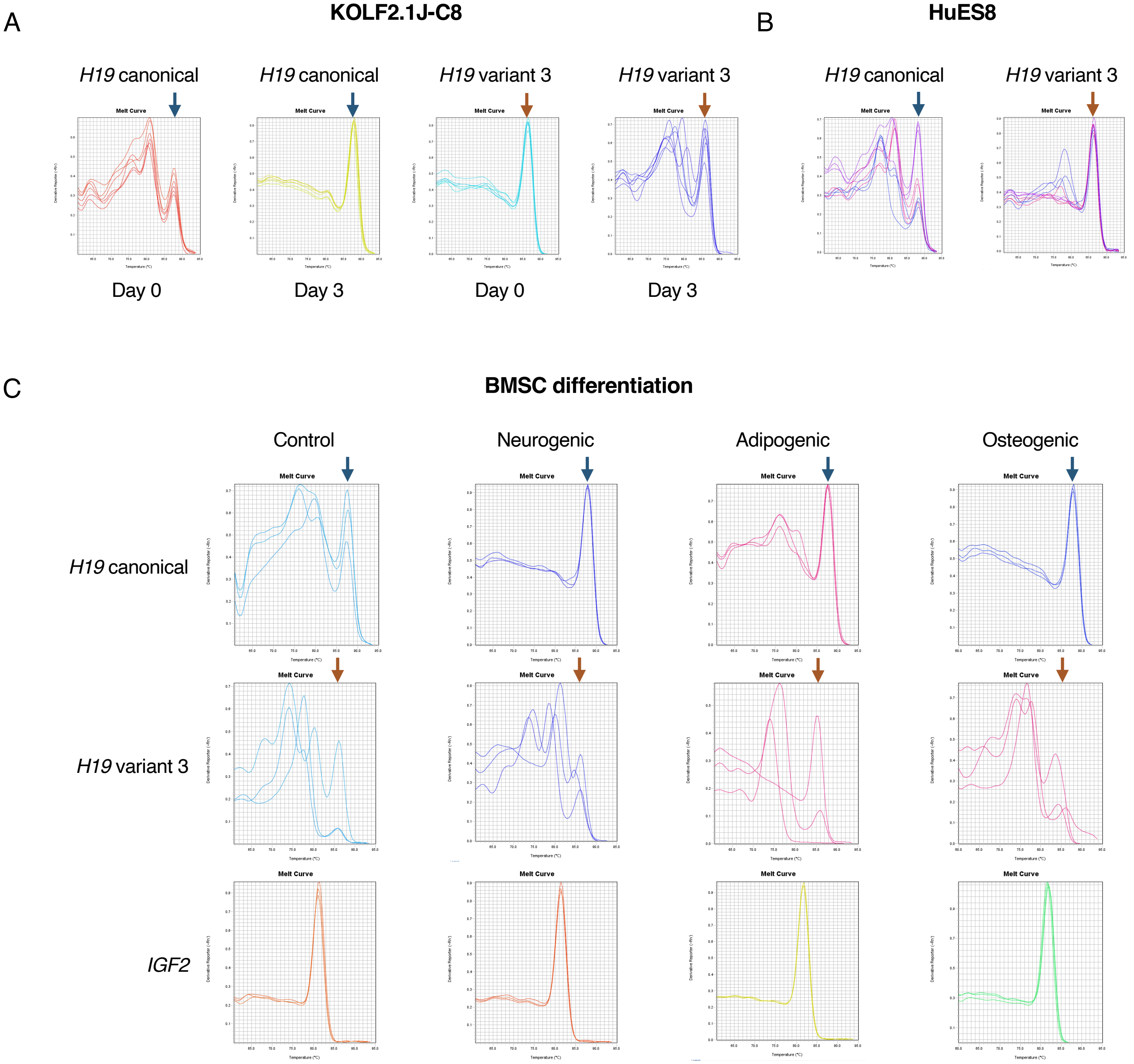


**Figure S2. Representative SYBR Green RT-qPCR melt curves during induced differentiation.** **(A)** Canonical *H19* and variant 3 assays in KOLF2.1J-C8 cells at days 0 and 3 of iNeuron differentiation. **(B)** Canonical *H19* and variant 3 assays in undifferentiated HuES8 hESCs. **(C)** Canonical *H19*, variant 3, and *IGF2* assays in control BMSCs and day-14 neurogenic, adipogenic, and osteogenic BMSC cultures. Blue and orange arrows mark canonical *H19* and variant 3-specific peaks, respectively, where present.
