## Supplemental Figure S3 for "*H19* isoform switching during neuronal differentiation reveals a distinct pluripotency-associated non-canonical transcript"

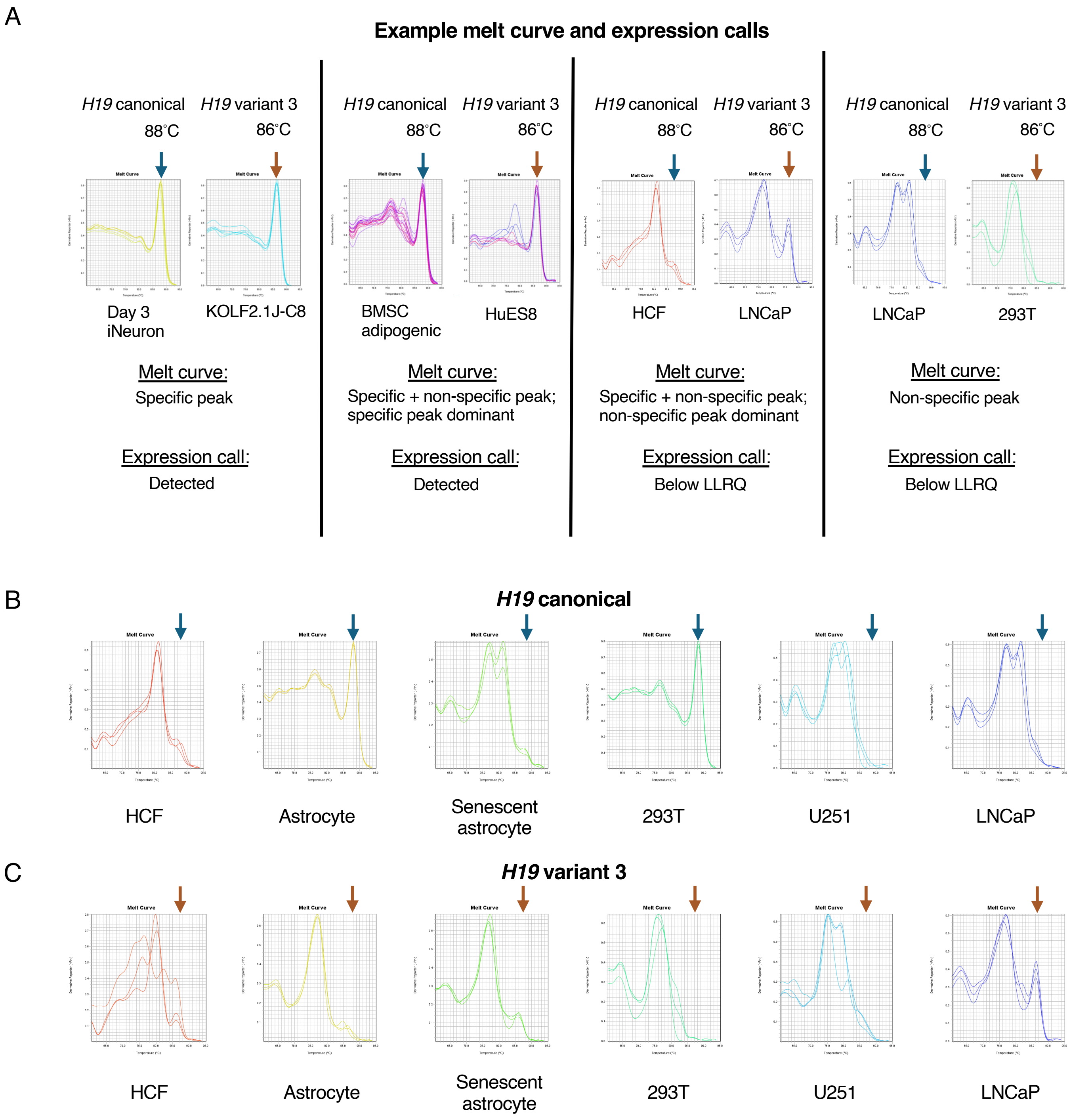


**Figure S3. Representative melt-curve profiles supporting canonical *H19* and variant 3 expression calls.**
**(A)** Examples of specific peaks, mixed specific/nonspecific profiles with either peak dominant, and nonspecific peaks only. “Detected” indicates a specific peak alone or dominant over nonspecific products. “Below LLRQ” (lower limit of reliable quantification) indicates that the specific peak is absent or does not dominate; this operational specificity-based classification is not a numerically validated quantification limit and does not establish transcript absence. **(B,C)** Representative profiles across additional cell types for **(B)** canonical *H19* and **(C)** *H19* variant 3. Blue and orange arrows mark canonical *H19* and variant 3-specific peaks, respectively, where present.
