## Supplemental Figure S4 for "*H19* isoform switching during neuronal differentiation reveals a distinct pluripotency-associated non-canonical transcript"

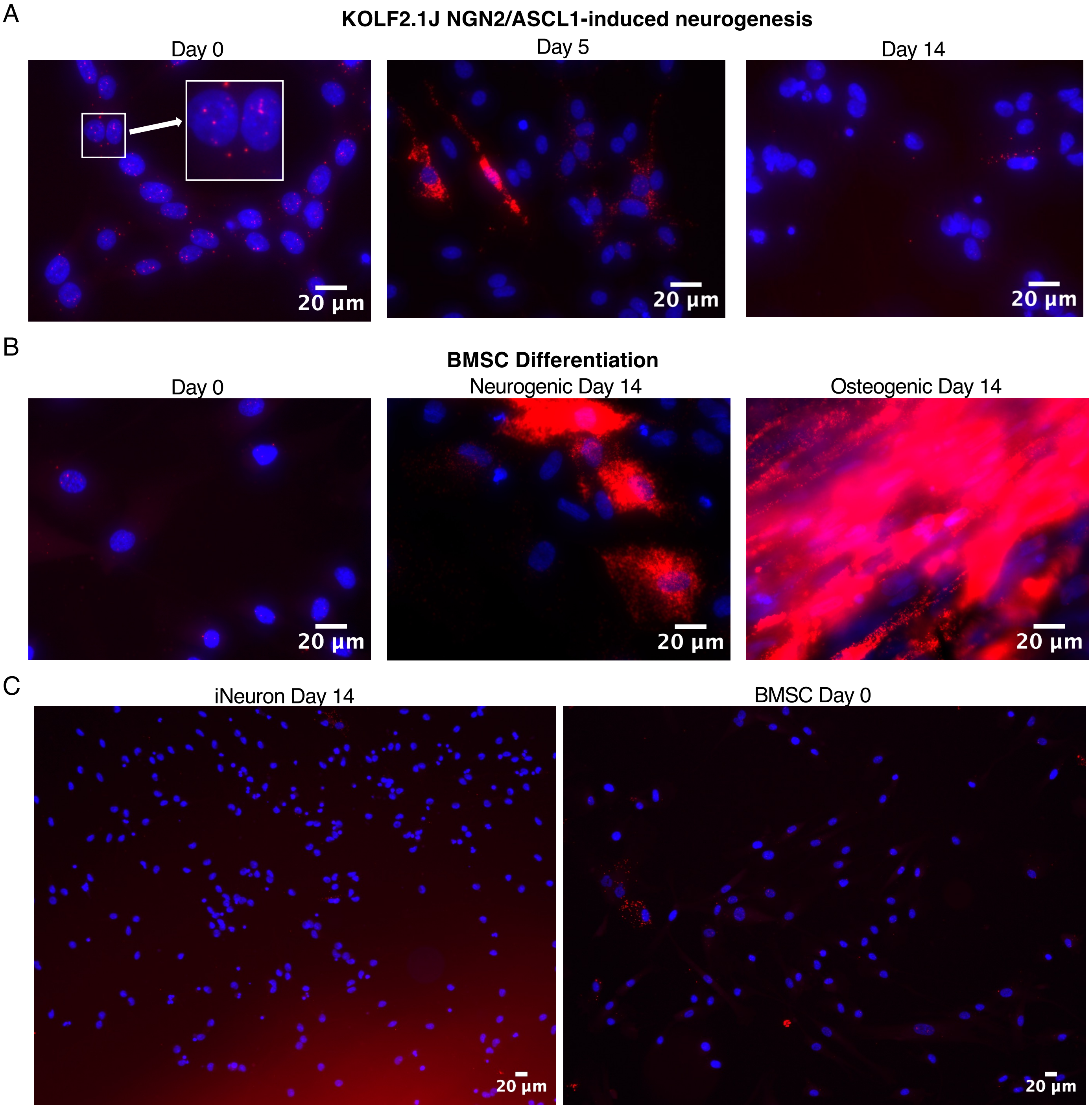


**Figure S4. Isoform-nonspecific *H19* RNAscope during NGN2/ASCL1-induced neurogenesis and BMSC differentiation.**

Representative *H19* RNA signal (red) and DAPI counterstain (blue) in **(A)** KOLF2.1J-C8 cells at days 0, 5, and 14 of iNeuron differentiation, with an enlarged day-0 region shown as an inset; **(B)** day-0 BMSCs and day-14 neurogenic and osteogenic BMSC cultures; and **(C)** lower-magnification day-14 iNeurons and control BMSCs. The probe recognizes sequences shared among *H19* transcript isoforms. Display brightness was increased by 50% in PowerPoint for all images except the neurogenic and osteogenic BMSC images in panel B. Scale bars as indicated.
