## Supplemental Figure S5 for "*H19* isoform switching during neuronal differentiation reveals a distinct pluripotency-associated non-canonical transcript"

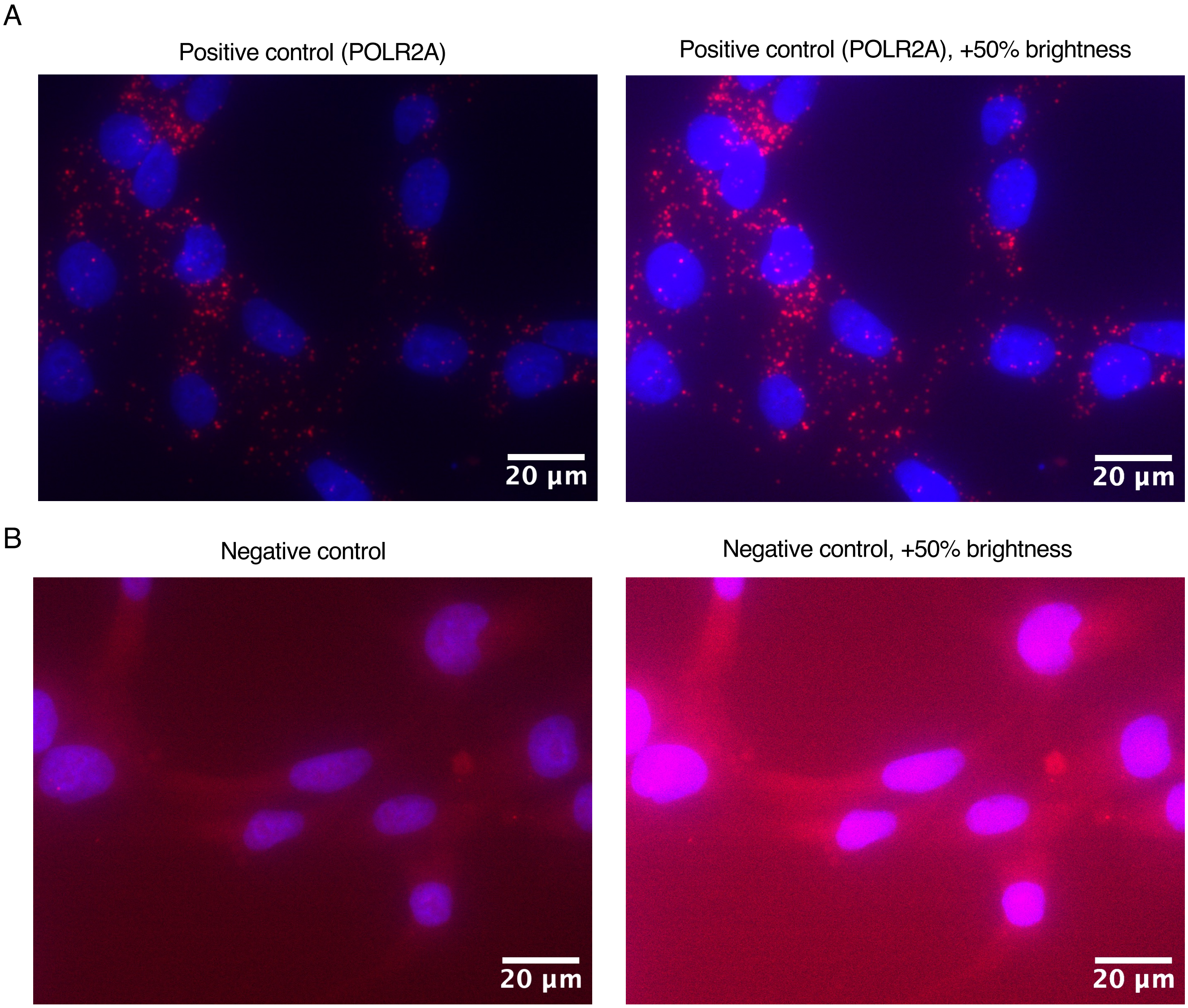


**Figure S5. RNAscope assay positive and negative controls.**
**(A)** Representative POLR2A positive-control and **(B)** negative-control probe images of KOLF2.1J-C8 cells. Each acquisition is shown at its original display brightness and with brightness increased by 50% in PowerPoint. Probe-channel fluorescence is red and DAPI is blue. The negative control shows diffuse red-channel background without discrete probe-associated puncta. Scale bars as indicated.
