## Supplemental Figure S6 for "*H19* isoform switching during neuronal differentiation reveals a distinct pluripotency-associated non-canonical transcript"

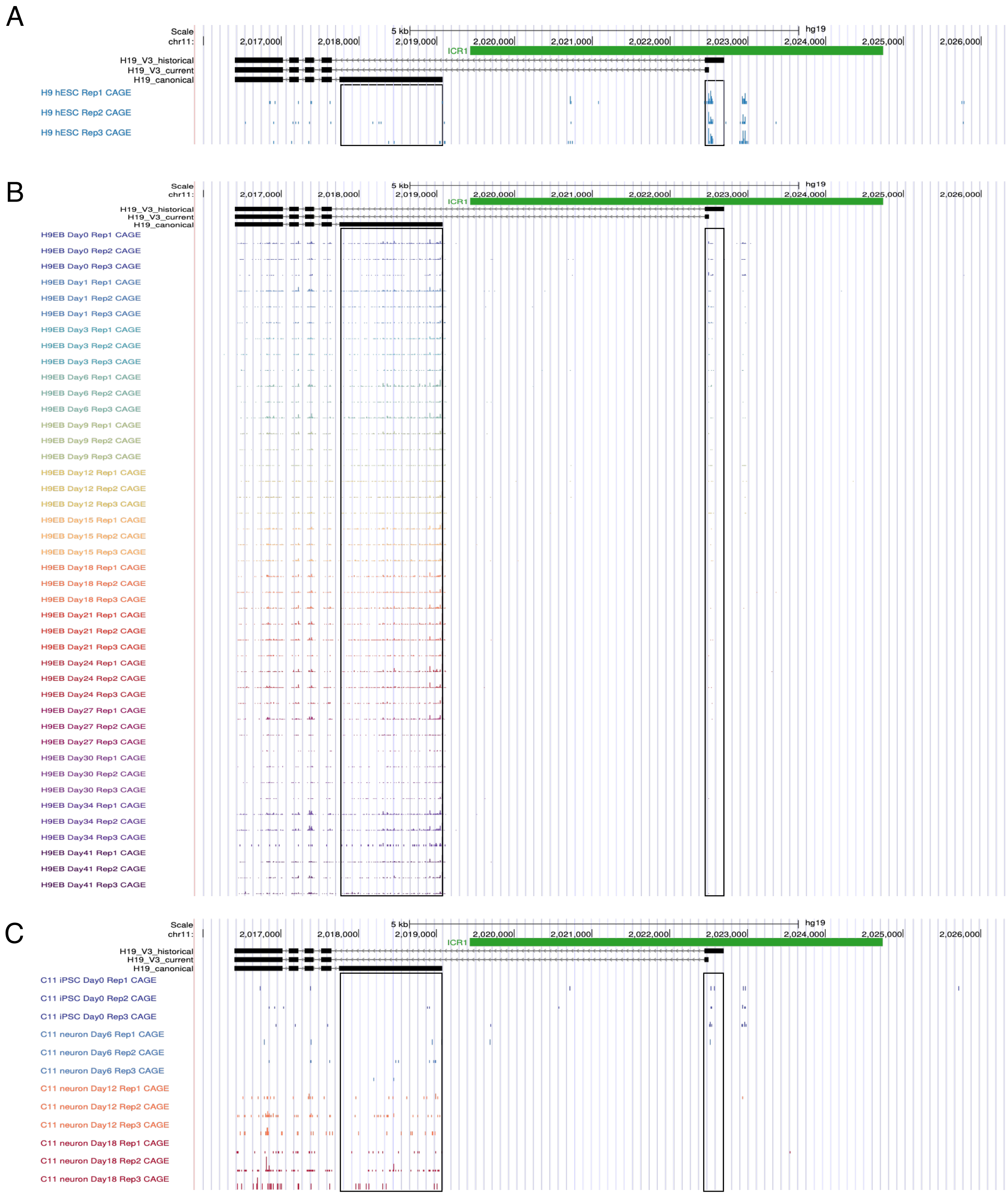


**Figure S6. Base-resolved FANTOM5 CAGE signal across the *H19* locus in pluripotent cells and during differentiation.**

Reverse-strand CAGE transcription start site (CTSS) tracks are shown for **(A)** H9 hESCs, **(B)** H9-derived embryoid body-to-melanocyte differentiation, and **(C)** C11 hiPSC-to-neuron differentiation. Each track represents one deposited sample. Boxes mark the canonical and historical variant 3 first-exon intervals quantified in Fig. 4. Canonical *H19*, current variant 3, historical variant 3, and ICR1 annotations are shown for orientation.
