## Supplemental Figure S7 for "*H19* isoform switching during neuronal differentiation reveals a distinct pluripotency-associated non-canonical transcript"

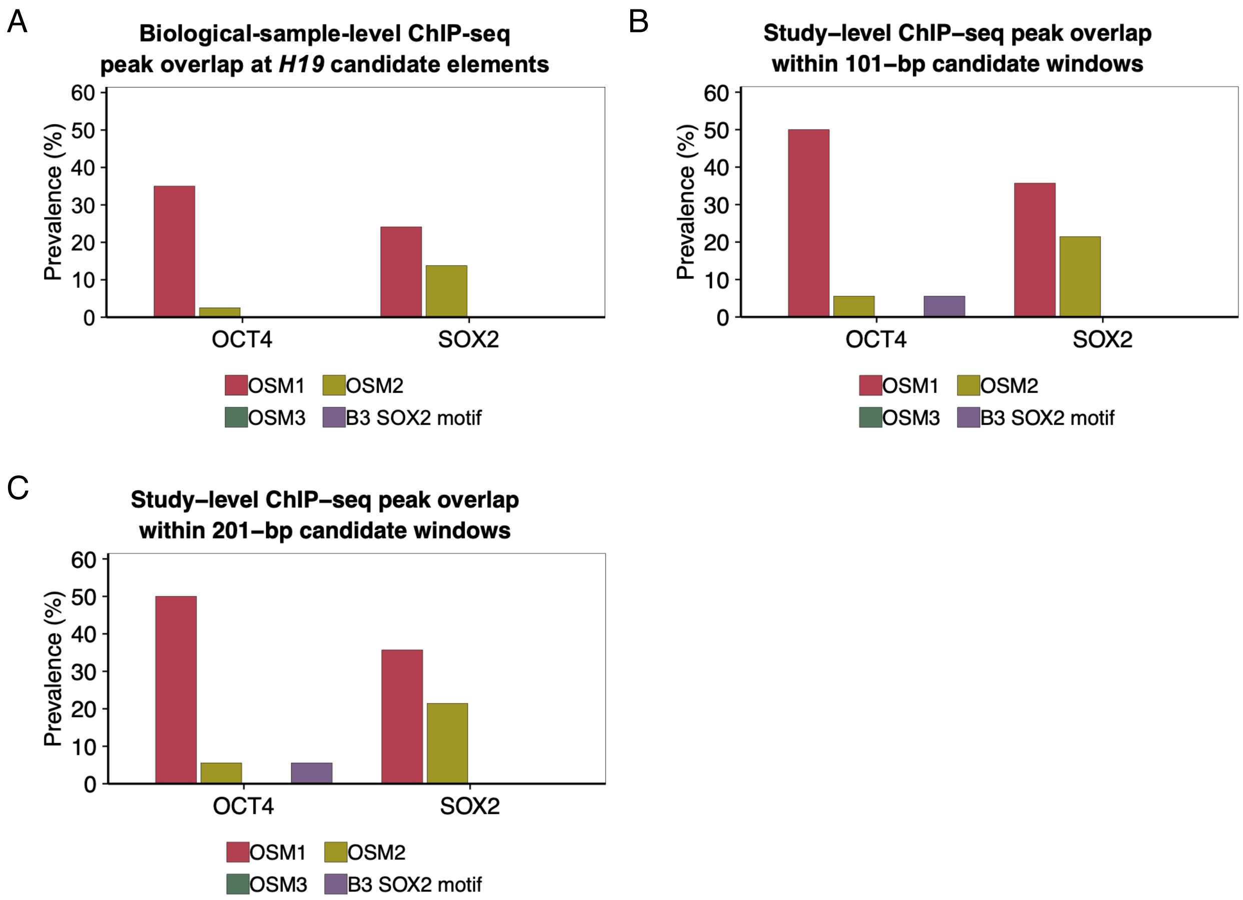


**Figure S7. Biological-sample-level and expanded-window sensitivity analyses of OCT4 and SOX2 ChIP-seq peak overlap at *H19* candidate elements.**

**(A)** Biological-sample-level prevalence of MACS2 peaks directly overlapping OSM1–3 or the B3 SOX2 motif after technical-replicate collapse. **(B,C)** Study-level prevalence using uniform **(B)** 101-bp or **(C)** 201-bp windows centered on each candidate element. A study was positive if at least one eligible biological sample contained an overlapping peak.
